# Sensing Local Field Potentials with a Directional and Scalable Depth Array: the DISC electrode array

**DOI:** 10.1101/2021.09.20.460996

**Authors:** Amada M. Abrego, Wasif Khan, Christopher E. Wright, M. Rabiul Islam, Mohammad H. Ghajar, Xiaokang Bai, Nitin Tandon, John P. Seymour

**Affiliations:** Department of Neurosurgery, University of Texas Health Science Center, Houston, TX, 77030 USA; Department of Bioengineering, Rice University, Houston, TX, 77030, USA; Department of Electrical and Computer Engineering, Rice University, Houston, TX, 77030, USA

## Abstract

A variety of electrophysiology tools are available to the neurosurgeon for diagnosis, functional therapy, and neural prosthetics. However, no tool can currently address these three critical needs: (i) access to all cortical regions in a minimally invasive manner; (ii) recordings with microscale, mesoscale, and macroscale resolutions simultaneously; and (iii) access to spatially distant multiple brain regions that constitute distributed cognitive networks. We present a novel device for recording local field potentials (LFPs) with the form factor of a stereo-electroencephalographic electrode but combined with radially positioned microelectrodes and using the lead body to shield LFP sources, enabling *<u>di</u>*rectional sensitivity and *<u>sc</u>*alability, referred to as the DISC array. As predicted by our electro-quasistatic models, DISC demonstrated significantly improved signal-to-noise ratio, directional sensitivity, and decoding accuracy from rat barrel cortex recordings during whisker stimulation. Critically, DISC demonstrated equivalent fidelity to conventional electrodes at the macroscale and uniquely, revealed stereoscopic information about current source density. Directional sensitivity of LFPs may significantly improve brain-computer interfaces and many diagnostic procedures, including epilepsy foci detection and deep brain targeting.

## 1. Introduction

When recording neural activity, what spatial resolution is required to elucidate the etiology or diagnosis of a disease? What time scale will best help understand the neurobiology of a given circuit? The question of scale is an ongoing challenge and our enthusiasm for more data is balanced by practical questions of safety and shortcomings of the recording technology. Non-invasive field potentials (electroencephalograms (EEG) and magnetoencephalograms (MEG)) are used to measure electrical activity within the brain but have limited spatial precision and requires activity from millions of neurons. Subdural surface electrodes (a field potential known as electrocorticogram (ECoG)) and/or depth electrodes are used instead of EEG/MEG when greater spatial resolution is required ^1^. Recently, it has become common for patients with drug-resistant epilepsy and likely focal seizures to undergo recording to localize the epileptogenic zone. These procedures were once based on surface arrays (i.e., ECoG) but have been largely replaced with stereo-electroencephalogram (sEEG) ^2^ sEEG is a specific form of a depth array with eight to sixteen ring electrodes and are usually only 0.8 mm in diameter, which is much smaller than other depth devices like deep brain stimulators. Given modern pre-op imaging tools and frameless robotic guidance, its advantages over the ECoG grid are a reduced risk of hemorrhage, infection, and migraines ^3–8^. A potential factor is their small access holes minimize damage to the highly vascularized skull and meninges. These advantages have been impactful, for example, in the diagnosis of medial temporal lobe epilepsy ^9^. In short, stereotactically placed depth electrode arrays possess an excellent form factor for minimally invasive large-scale recording and stimulation, and concurrently enable an interface with broadly distributed cortical networks.

The success of depth arrays implies their use in more applications such as a brain-computer interface (BCI), although the decoding performance of the sEEG ^10, 11^ is still trailing that of surface grids ^12–14^ and the Utah array ^15, 16^. *The amplitude and spatial resolution of sEEG recordings in their current form is limited by the circumferential nature of the ring electrode as we will discuss in detail*.

Besides field potentials (FPs), researchers have two other temporally precise recording domains: extracellular single cell (300-5000 Hz) and local field potentials (LFP, < 300 Hz). The former is generally preferred in systems neuroscience and BCI but measure cellular action potentials only within 100-120 μm of the source ^17^. This limits the volume of cortex sampled in the high-frequency band, but microelectrodes can simultaneously record the LFP over large distances^18^. A common misconception about microelectrodes is that they have high local sensitivity and poor “macro” sensitivity relative to a ring electrode; however, it records everything a macroelectrode records and more. By the law of superposition, it will sense every distant field potential and the slow voltage changes of local synaptic activity as well as the higher band spikes. LFPs when measured from microelectrode arrays with little insulating substrate (microwires, Utah array, or any thin-film probe^19, 20^) are data rich, but without source separation the true information capacity is low. To improve the information capacity of LFP recordings, therefore, the goal is to measure fewer LFP sources and with greater fidelity until source separation is achievable.

We describe a novel multi-scale recording array capable of stereo-LFP and current source density measurements. The *di*rectional and *sc*alable electrode array (DISC), designed to overcome the limitations of signal fidelity and source separation, provides much needed information at the mesoscale (0.1-5 mm). We performed finite element modelling based on electro-quasistatic physics and *in vivo* recording in the rat barrel cortex. An under-appreciated phenomenon, “substrate shielding”, was applied to spatially segregate LFPs. Others before us were the first to identify the phenomenon but on a smaller scale. Two reports ^21, 22^, and later others and ourselves ^23^ describe how neurons can be shielded by the device, resulting in directional sensitivity. Boston Scientific has also begun offering directional electrodes in the form of a ring with 3 segmented sections across a 1.2 mm diameter DBS lead body. Recently, detailed modeling by Noor and McIntyre ^24^ describes the quantitative advantages of the segmented electrodes versus ring electrodes in providing directionality. Our research expands these concepts in a more generalizable design and establishes why microelectrodes on larger substrates may be transformative to neuroscience and clinical diagnosis. We quantify how source separation may be affected by substrate diameter, electrode size, electrode configuration, and source orientation and distance. In addition, we demonstrate directional accuracy and multidirectional current source density analysis with histology. DISC provides low-noise microscale or macro-equivalent recordings, which suggests clinically available sEEG electrodes can be imitated when replaced by DISC. Finally, we describe several manufacturing approaches for making DISC in a way that is compatible with the goal of being concurrently useful as a standard-of-care stereotactic electrode.

## 2. Results

### Electro-quasistatic model of substrate shielding and directionality

Maxwell’s equations can be reduced to an electro-quasistatic model with the current conservation law applied to an array of sinks and sources in the extracellular space ^25, 26^. These results apply to all frequencies of clinically relevant extracellular potentials. The Poisson equation,

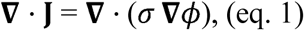

where J is current density, σ is conductivity, and represents the extracellular electric potential, can be applied to arbitrary current sources in a conducting volume with boundary conditions including the insulating body of a recording device.

#### Directionality as a function of substrate diameter

We compare a narrow substrate (e.g., microwire or Michigan style array), a ring electrode, and DISC in the presence of a dipole approximating a cortical column from layer V pyramidal cells (1.2 mm height and 2.0 nA*m/m^2^ current dipole moment) ^27^ (Fig. 1). A useful measure for the “directionality” of a source and device combination is the change in voltage between nearby electrodes, such as the front voltage to back voltage ratio (F/B). As shown, the F/B ratio has an exponential relationship with substrate diameter, which has not been previously reported. Large diameter insulating bodies create a disturbance in the voltage contours unlike smaller diameter recording tools. Since this demonstration creates an increasing distance between electrode pairs, it is important to compare this ratio when lead body conductivity matches the value of tissue (σ = 0.2 S/m) (Fig. 1C green line). The difference between these lines represents the effect of “substrate shielding” on the dipole current, which is caused by diverting the current around the lead body (Suppl Fig. 1). The isopotential around the front electrode extends toward the source, which is the cause of the absolute voltage amplification (Fig. 1C (dashed line)). Similarly, the isopotential around the back electrode extends away from the source because current is not flowing in the immediate vicinity (Extended Data Fig. 1). These combined effects provide a previously undescribed directionality available to a larger diameter device when recording field potentials even at considerable distances.

**Figure 1.**
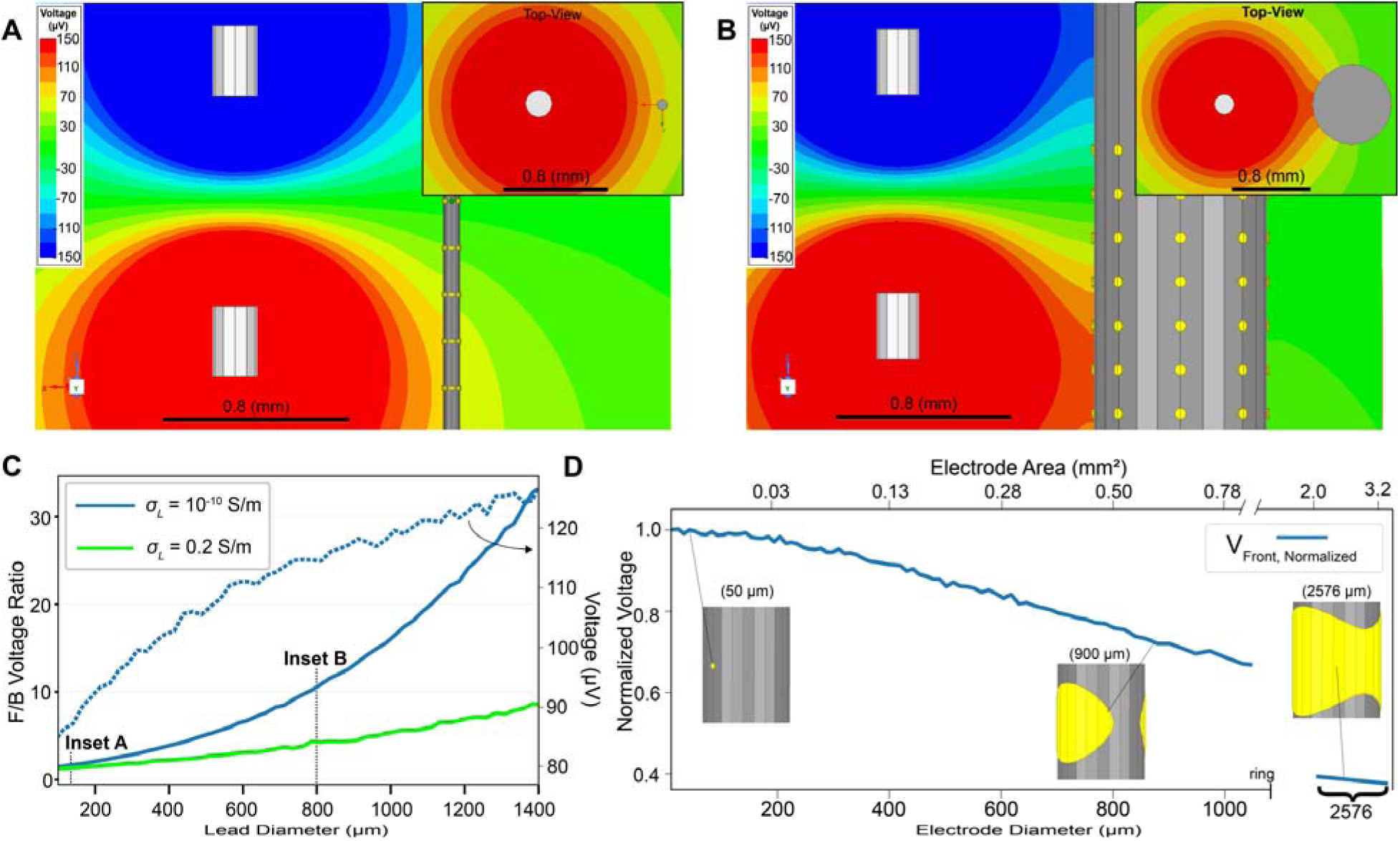
Directionality and amplitude as a function of substrate and electrode diameters. **A,** A 65 μm diameter and **B,** 800 μm diameter substrate with independent electrodes on opposite sides at a fixed distance from a dipole source. Voltage V from a current source shown to left of the lead body represents activity in layer V pyramidal cells in the rat cortex (Ø: 200 μm) generated in a finite element model (FEM). Voltage is inversely related to distance and perturbed by changes in conductivity σ in space created by the insulating body. **C,** FEM model (ANSYS) of the front to back electrode voltage ratio as a function of diameter (Ø_sh_). When σ of the lead body matches local tissue (σ = 0.2 S/m), F/B increases due to the increasing distance between front and back electrodes. Substrate shielding magnifies the difference between the front and back electrodes and amplifies the voltage at the front electrode (dotted trace). **D,** Normalized voltage amplitude as a function of electrode diameter using the same dipole source. Attenuation becomes significant beyond 120 μm. A ring forms at 1238 μm (front and back merge) and the amplitude is attenuated by 60% relative to baseline. Increasing the ring diameter any further has negligible attenuation. A video of this effect is provided in Suppl. Movie 1.

#### Electrode diameter and LFP source amplitude

Large electrode surface areas are most practical in noisy clinical environments given their low impedance and therefore poor electromagnetic interference (EMI) pickup. We parameterized the electrode diameter on an 800-μm lead body and measure the peak amplitude when a dipole and the nearest electrode had a gap of 0.8 mm. The decline in amplitude with increasing diameter is quite significant even before the electrode wraps around the substrate to form a ring (Fig. 1D). Once the ring forms, the area no longer has any significant affect. A 120-μm diameter microelectrode (11,300 μm^2^) has only a theoretical loss of amplitude of 3% for dipole-equivalent LFP sources. A minor attenuation may be an excellent tradeoff given some degree of EMI and thermal noise immunity, and therefore achieve greater SNR. Clinically available directional leads have a diameter of 1.2 mm (DB-2202, Boston Scientific). The FEM model predicts that even a 1-mm diameter electrode will be attenuated by 27% relative to even a 120-μm electrode. Given that the actual electrode is a 1.5-mm tall rectangle, the attenuation is likely greater.

#### Source geometry and orientation

The model so far considered an orientation most likely encountered when the array implantation is normal to the neocortical surface and parallel to cortical columns. Many device trajectories ^28, 29^ and therefore dipole orientations are possible. Open field dipoles may also appear as local sinks or sources (monopoles) and represent the best-case attenuation (∼1/r). Each of these situations can create degrees of directionality and attenuation profiles. We analyzed several possibilities: a monopole (200 μm), dipole (200 μm), large dipole (800 μm), and orthogonal dipole all at varying distances from 0.15 mm to 5 mm, defined as the mesoscale, by Nunez et al. ^30^. The F/B ratio over distance for each is a useful quantitative juxtaposition. As expected, a parallel-oriented dipole has the greatest directionality but the least range (falls to 20 μV at 2.1 mm). The orthogonal dipole, same diameter, has a flatter falloff with a 20 μV distance equal to 2.7 mm). Larger dipoles (Ø: 800μm) can easily extend beyond 4 mm (Suppl. Fig. 1, Suppl. Movie 2).

#### Directionality in a noisy multi-source environment

In another model, we introduced eight cortical sources within ∼5 mm of the device (Fig. 2), added independent, gaussian-distributed noise to each electrode, and measured the SNR during an evoked potential task to isolate each source based on phase (see *Methods*). Our simulation provides a challenging scenario by varying distance and diameter dramatically (Fig. 2A, B, Suppl. Table 1). Source 1 for example has two other sources within 20 degrees and therefore communicates over the same column of electrodes. Similarly, source 3 and 4, and 5 and 6 have similar angles. Source 1 at 1 mm distance has an SNR = 1.8 dB with no repeated trials and 7.6 dB at 50 trials (Fig. 2C). For an ultra-high-resolution ring electrode (0.4-mm tall, 0.8-mm diameter) the SNR was 0.38 and 3.9 dB. DISC microelectrodes were also assigned 4.3 μV_RMS_ (root mean square), whereas the ring electrode included only 2.7 μV_RMS_. DISC does not usually have lower noise (data not shown) but does have greater SNR for each of the 8 sources. Distant sources 4, 6, and 8 had relatively modest improvement using DISC, but the larger source 7 created a particularly large SNR relative to the narrow substrate or ring electrode. Common average referencing (CAR) improves SNR for the farthest sources by subtracting out biological background noise. The ring electrode outperforms a microelectrode on a small substrate (e.g., microwire) for all but the closest source because of its lower noise floor. A 2-mm tall ring electrode was added for completeness and its large size reduced the amplitude for all but two sources (Suppl. Fig. 2). Arrays of microelectrodes would have also benefited from CAR ^31^ although it was not tested here. An additional model of tissue encapsulation conductivity and thickness was performed (Suppl. Fig. 3). Encapsulation tissue improved the recording amplitude slightly as predicted in Moffitt, et al. ^22^.

**Figure 2.**
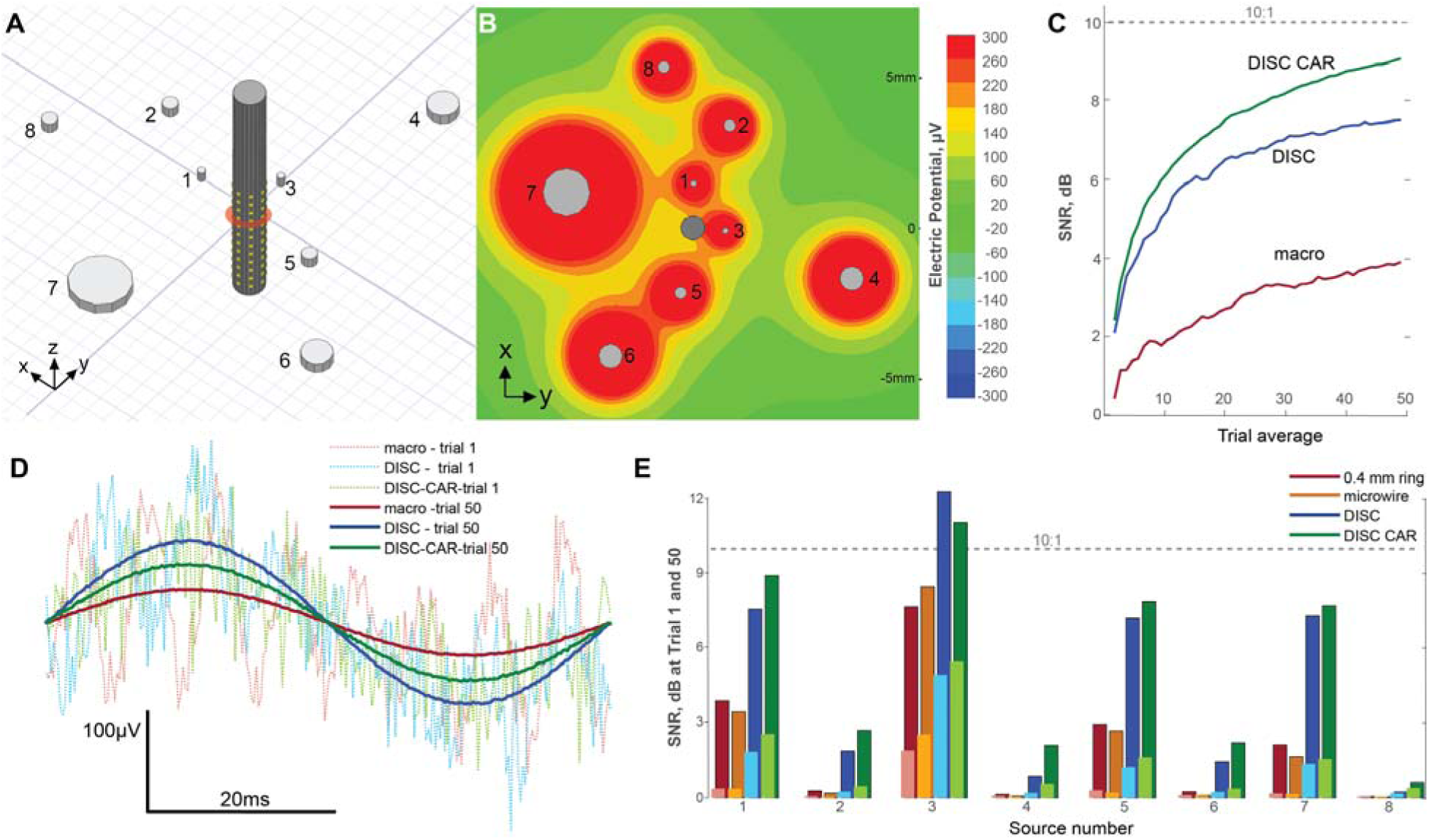
Electro-quasistatic 3D dipole model demonstrating directional sensitivity in a multi-source configuration. **A,** 8 simultaneous dipoles modeled in FEM with identical surface boundary current density (only dipole sink is shown, 0.5-mm grid). Three device types are modeled: DISC (shown), microwire, and 0.4-mm tall ring electrode. **B,** Voltage heat map through layer V dipoles with sources on at peak current density (1.67 μA/mm^2^). Heat map also intersecting the dipole sinks. **C,** Signal-to-noise (dB) for macro, DISC, and DISC with CAR during trial 1 and cumulative trials. **D,** Waveform examples for 1 and 50 trials for the macro and DISC electrode when phase locked to Source 1. Sources 2-8 are assigned a random phase and frequency. Assigned noise of 2.7 μV_RMS_ and 4.3μV_RMS_ to each ring or microelectrode, respectively. **E,** SNR comparison of the simulated potentials for each source independently phase-locked. Microwire Ø=65μm. Trial 1 = light color; Trial 50 = dark color (avg).

Virtual electrode geometries can be computed with digital averaging and referencing ^32^. Generating an equivalent macroelectrode signal is important because of its long clinical history and effective filtering in certain frequencies ^33^. We validate this ability both theoretically and empirically. First, we used ANSYS and the previously described multi-source and noise simulation model. For each of the 8 sources in our model, we measured both the ring and virtual ring (comprised of 3 x 8 electrodes at the same depth) and found DISC performed nearly equivalently across sources (Extended Data Fig. 2). Second, we demonstrate that the average noise value *in vivo* on a virtual 0.4-mm and 2-mm ring electrode was 3.9 and 3.1 μV_RMS_, respectively (Suppl. Table 2).

### *In vivo* amplitude, SNR, and directional sensitivity

#### Amplitude and SNR

One tetrode and DISC were implanted into 9 anesthetized rats and 9 whiskers were stimulated over 450 trials. We compared recordings from the tetrode, DISC, and two virtual ring electrodes created from DISC recordings. One ring type was a 0.4-mm tall x 3 array and the other a 2-mm tall single electrode (mirroring the commercial standard). The mean amplitude of the LFP signal of our DISC device (249.3 ± 183.5 µV) was significantly better than the tetrode (86.6 ± 65.5 µV, Tukey’s test p<0.001), virtual 0.4-mm ring (120.3 ± 97.5 µV, p<0.001), and the virtual 2-mm ring (59.2 ± 63.3 µV, p<0.001). The DISC device also significantly improved the LFP SNR (DISC: 13 ± 3.3, tetrode: 7.9 ± 3.3, virtual 0.4-mm ring: 10.3 ± 3.5, virtual 2-mm ring: 7.3 ± 4.0 dB p<0.001) (Fig. 3). Suppl. Table 2 contains the P-P and RMS amplitudes and noise floor values (100 ms window prior to stimulus) for all devices, including DISC with CAR. The amplification of DISC electrode amplitude (P-P) relative to the tetrode was 214% (relative median values). The amplification of the source is putatively a combination of substrate amplification (Fig. 1C) and placing electrodes closer to the source (Suppl. Fig. 4).

**Figure 3.**
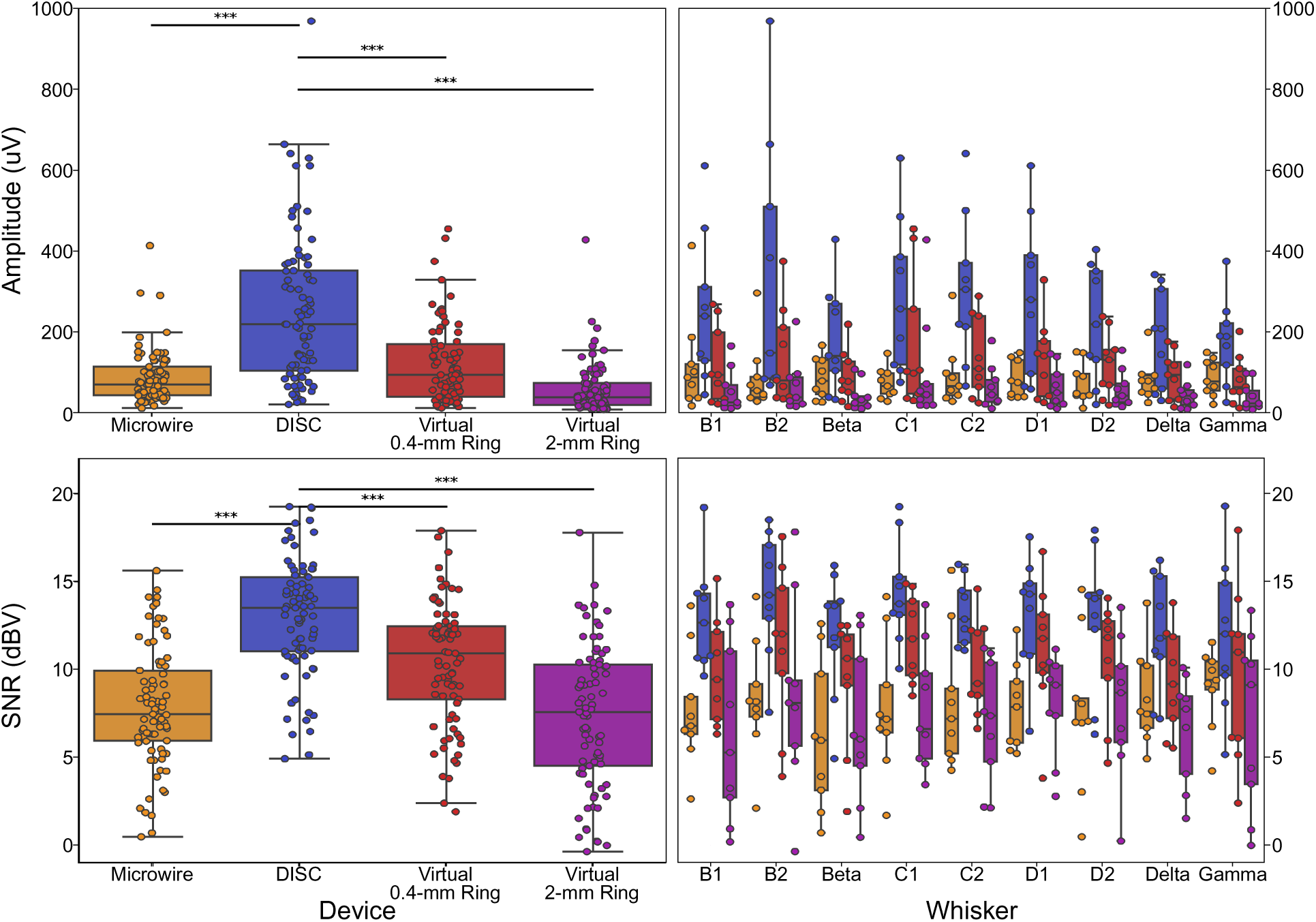
Amplitude and SNR for each device type. (N=9 subjects and 9 whisker values each.) Ring electrodes were created virtually by averaging an array of DISC channels forming a virtual ring of height 0.4 mm or 2 mm. Vertical pitch of DISC array was 200 μm. Whisker by whisker values are shown to the right (N=9).

In practice, CAR improved the SNR in trial averages, increasing it by 2 to 15.3 ± 2.7 dB (Tukey’s test, p<0.001). A lower amplitude was also noted for CAR (Suppl. Fig. 5), as expected from theory. All experiments were recorded in a Faraday cage. To illustrate noise immunity with CAR, we also recorded a session with an intentional ground loop and no Faraday cage, which in this example made a significant, observable difference (Extended Data Fig. 3).

#### Directional Sensitivity and CSD

Cytochrome staining was performed in subjects S6, S8, and S9 to identify the device location. Histology from subject S6 indicated DISC and the microwire array were implanted in barrel E1 (medial edge of barrel field). A directional voltage profile (detailed in *Methods*) for nine whisker stimuli also correlated to the position identified in histology (Fig. 4, Suppl. Fig. 6, 7). We also modeled dipoles at the specific histological coordinates and diameter for each of nine barrels and plotted the FEM predicted profiles (Fig. 4C). Our model assumed a perfectly confined excitation pattern instead of the expected lateral excitation between barrels ^34, 35^, which would at least partially explain the center-focused profiles found *in vivo* relative to FEM. Directional profiles showed greater variance from trial to trial, as expected given independent electrode noise (Suppl. Fig. 8).

**Figure 4.**
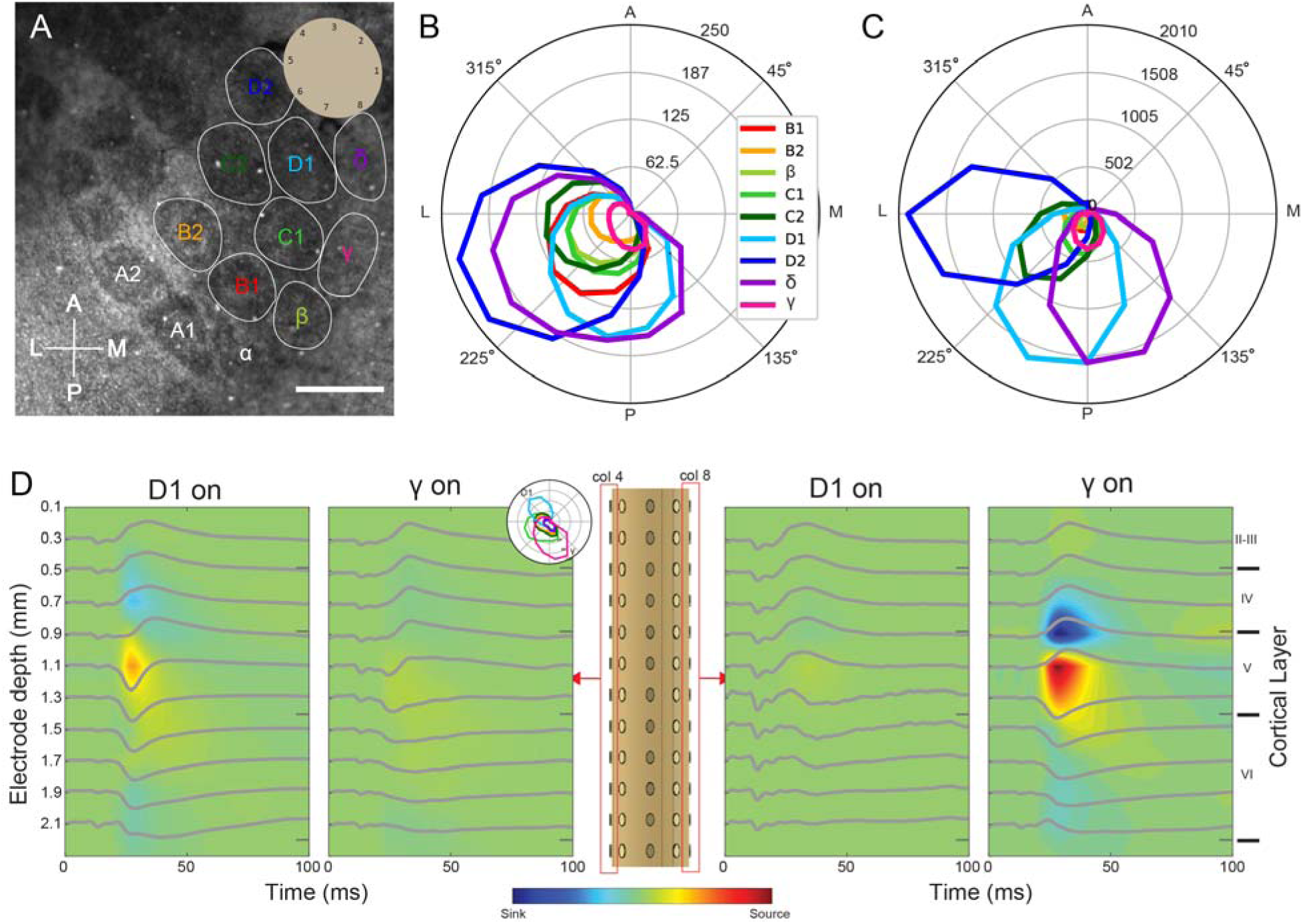
Demonstrating the directional sensitivity in rat barrel cortex. **A,** Example histology results using subject S6 and electrode layout from device 3. Unedited histology in Suppl. Fig. 6. Scale = 500 μm. **B,** LFP data from subject S6 and nine whiskers (row 6). Directional polar plot formed from 8 channels, interpolated to 16 channels (∼22.5°), and subtracted the minimum amplitude. **C,** FEM results assuming nine dipole sources spaced in an identical location as shown in the cytochrome-stained image. **D,** Multi-directional CSD from DISC demonstrated in subject S7 when two independent primary whiskers are stimulated. Barrel D1 was located closest to device column 4 while barrel γ was located on column 8 (∼180°). Polar plot of all whiskers shown in inset. Distinct CSD amplitude attenuation is observed. Average from 450 trials.

In subject S7 we performed CSD analysis to evaluate its ability to conduct stereo-CSD measurements. CSD is a solution to equation [1] given known potential values across the lamina ^36^. The results demonstrate CSD from two opposing direction in an example using barrels D1 and gamma (Fig. 4D).

### Whisker decoding and information capacity

Using the whisker stimulation experiment described above, we tested the accuracy of each device type in a 9-class discrimination task (Fig. 5), which is a measure of source separation. Our basic feature matrix prior to principal component analysis (PCA) consisted of the gamma band waveform for each included electrode. Each row represented one trial. PCA was applied for dimensionality reduction before classification. Linear discriminate analysis and 10-fold cross-validation was performed for all classifications.

**Figure 5.**
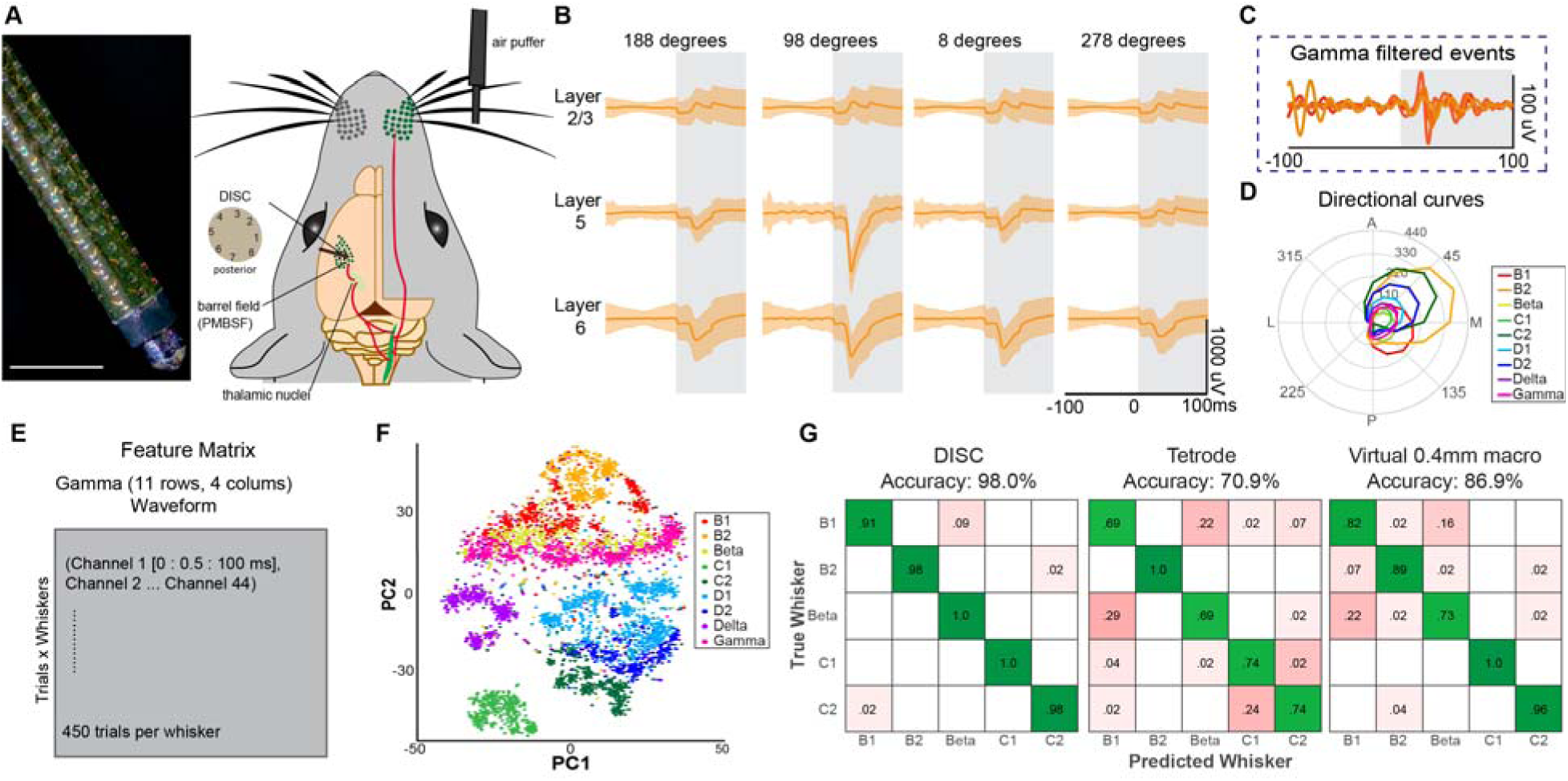
Methods and feature engineering overview for whisker stimulus discrimination in subject S4. **A,** Experimental design and DISC image. Scale = 1 mm. **B,** Example 3 x 4 arrangement demonstrating evoked response across four directions in whisker B2. **C,** Example of gamma filtered response of B2 at 98 degrees. Individual trials represented with different colors. **D,** Polar plot or directional curve calculated using the evoked response across eight directions for each whisker. Minimum value has been subtracted for contrast. **E,** Feature matrix arrangement. **F,** Dimensionality reduction and source separation using PCA. **G,** Example confusion matrix for three device types after analysis with a linear discriminant analysis (LDA) model.

DISC accuracy when using 11 rows and 4 columns was an average of 93.6 ± 7.7% and was significantly better than the tetrode, the 0.4-mm x 3 virtual ring array, and the 2-mm virtual ring (Fig. 6). The 3-ring virtual device performed second best and was significantly better than one 2- mm ring, which is the size of a standard sEEG. The classification accuracies of different electrode configurations and devices were tested and summarized in Extended Data Table 1. The best electrode configuration was 11 rows x 8 columns and 16 rows x 8 columns, but the 11 x 4 configuration was not significantly different and thus used in the device comparisons (Fig. 6).

**Figure 6.**
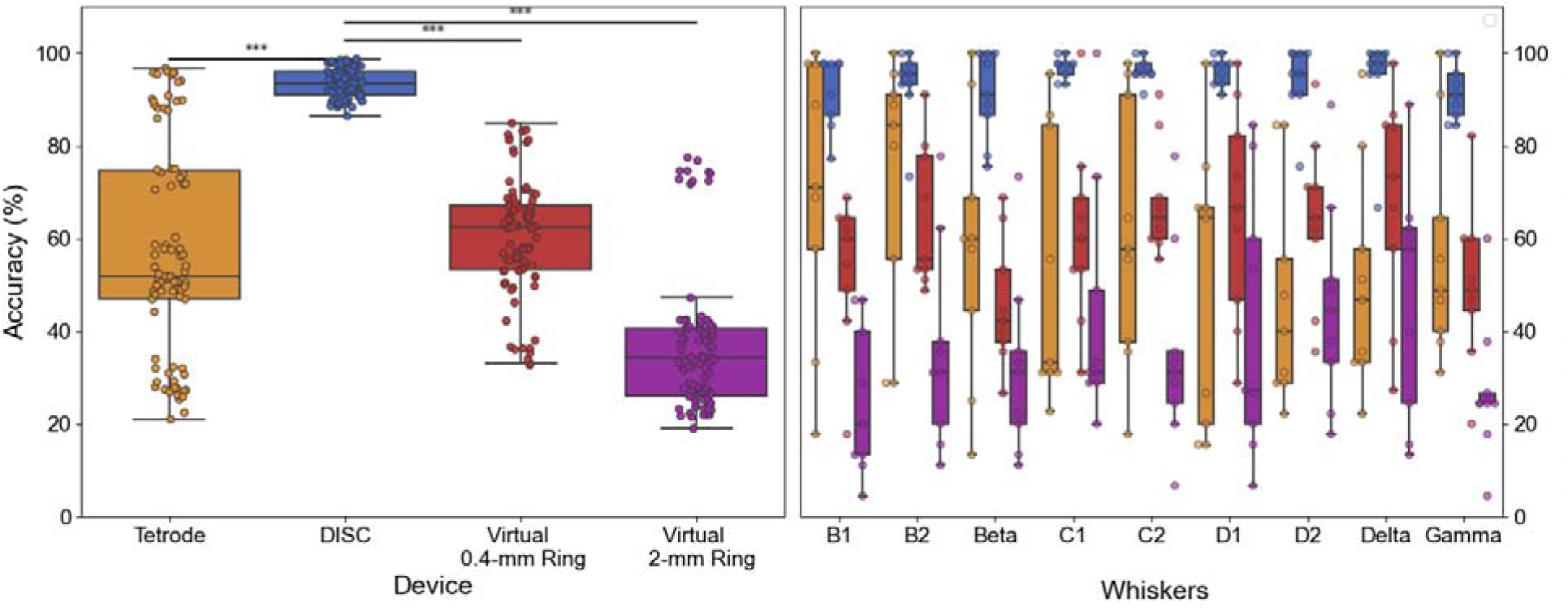
Decoding accuracy for each device type using a linear discriminant analysis model. (*** denotes p<0.001, generalized mixed effects model, 9 subjects, 10-fold cross-validation). Whisker by whisker values are shown to the right.

Overall, the electrode configuration provided the strongest influence on DISC accuracy. The 11 rows selected were in layers I to VI and the 4 columns were alternating to reduce redundancy. An important contrast is the DISC 1 x 4 configuration with the tetrode to remove the effect of channel count. We chose 4 orthogonal electrodes on DISC in the same layer as the implanted tetrodes (1.2 mm deep). DISC “tetrode” significantly outperformed the tetrode with an accuracy of 73.4 vs 58% (p<0.001). The amplification and substrate shielding effects can best explain this performance (Fig. 3). Further evidence of the impact of substrate shielding is the accuracy of a 11 x 1 (similar to a linear U-probe ^37^) vs a 1 x 8 DISC configuration, which was 68.8 vs 78.1%, respectively (Extended Data Table 1). Despite fewer channels, this single row input was more predictive than a linear array spanning the entire cortex.

### DISC manufacturing

A variety of manufacturing methods were tested and conceived to improve the likelihood of translation. Prototype linear arrays were manufactured in the cleanroom at Rice University using a polyimide-metal-polyimide process as previously described ^38^, but aligning 4 arrays per lead body resulted in poor alignment and a complicated backend design (Fig. 7A, backend not shown). Next we purchased similarly manufactured arrays from a commercial source originally designed for high-density ECoG (Diagnostic Biochips, Inc) and used an assembly technique previously described ^39^(Fig. 7B). The first challenging aspect of this method is the tight tolerance between the mold and device. Also, if epoxy is used, the needed tight tolerance creates large capillary forces and often coats electrodes during the curing process. Another assembly approach is wrapping the array using heat shrink (details in *Methods*). We patterned a thin (20 μm) adhesive sheet and mount to the backside of the 128-channel array. We prepared an insulated stainless-steel wire (432-μm diameter). Finally, we wrapped the components using heat shrink and heat gun (Fig. 7C). This provided the greatest electrode yield and un-altered impedance values. There are many future options including wrapping on a silicone cylinder, direct molding of the PI array, and additive manufacturing. We also modeled the stiffness of a thin-film, polyimide array wrapped over a silicone core, such as medical-grade tubing. The stiffness of a silicone-based DISC even with a polyimide wrap would be almost 1/100^th^ the stiffness of the rigid DISC used here, but it also would be 2.7X stiffer than a standard DBS lead body (Suppl. Fig. 9). Given the compression limits of heat shrink, and the stiffness of the final assembly, we recommend polyimide films be no thicker than 12 μm and as thin as 6 μm, below which kinking of the substrate is more likely.

**Figure 7.**
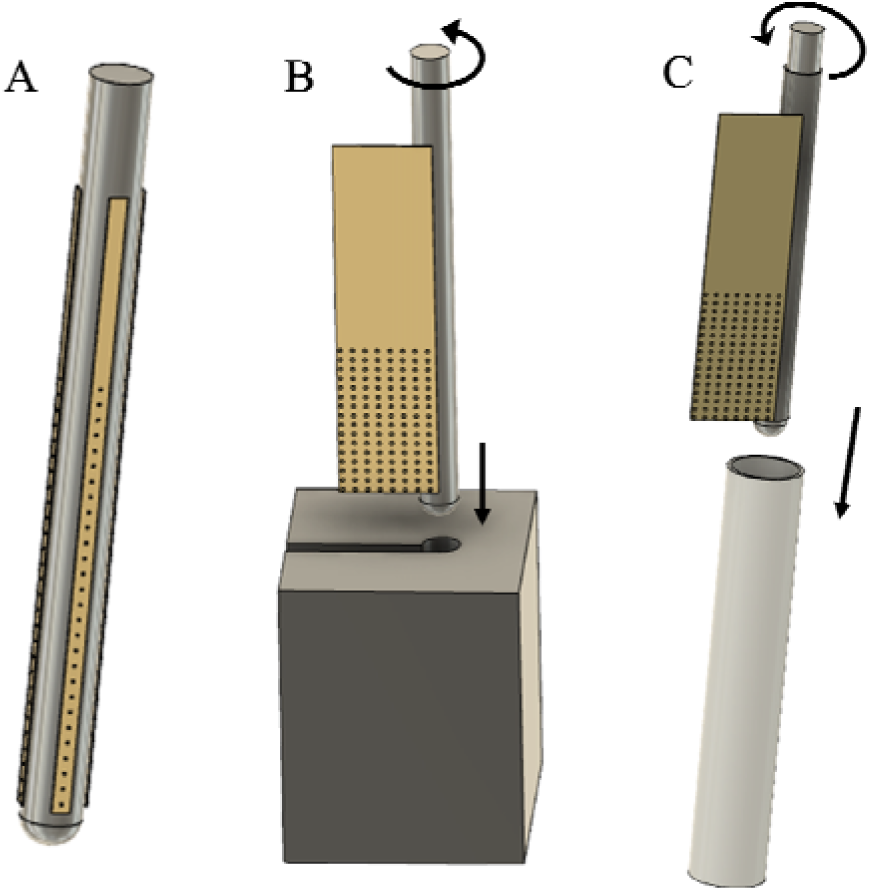
Example fabrication and assembly schematics available to practitioners. **A,** Linear polyimide arrays attached to rigid cylinder. **B,** Wrapping onto a rigid cylinder inside a rigid mold (originally proposed by ^39^). **C,** Wrapping method over a rigid body using heat shrink became our preferred method.

## Discussion

The *in vivo* demonstration agreed well with the predictions of the electro-quasistatic model, despite the potential for local damage of DISC relative to a much smaller tetrode. DISC was effective at the whisker classification task over the ∼1.4 mm tested, despite its relatively narrow footprint. In subject S6, the furthest barrel was 1.42 mm away (β). The accuracy of this subject remained very high despite the barrel distance (92-100%, 10-fold cross-validation, Suppl. Fig. 6). As predicted by the Shannon-Hartley theorem, higher SNR was related to greater information capacity (Fig. 3)^40^. This is supported by the virtual tetrode performance of DISC against the real tetrode comparing only 4 channels of information. The spatial diversity of DISC provides a unique advantage relative to the other tested devices and yet it did not require a large craniotomy required for a Utah array or an ECoG array. The vertical span and the directionality are the most likely factors enabling decoding success, although of those two, we demonstrated that a 1 x 8 (directionality) was more predictive than the 11 x 1 (vertical span). More electrodes resulted in higher accuracy (Extended Data Table 1), but these were relatively small gains – a simple 3 x 4 decoded at 89.3% accuracy whereas 11 x 4 was only 4.3% more. To identify the most discriminating features, 8 single statistics were selected and used for classification. Our conclusion from this analysis is that accurate source separation requires a diverse set of information including variations of amplitude, frequency, direction, and phase in whisker discrimination (Suppl. Table 3).

Our results demonstrated for the first time, as far as we are aware, multi-directional CSD recordings. This capability may, for example, contribute to the understanding of lateral excitation or inhibition between cortical columns ^41^. The insertion of DISC, given its diameter, may damage a portion of the local cortical column. In a recent histological study of a 800-μm diameter sEEG (N=3 devices), the measured radius of tissue damage in a non-human primate was only 50 μm for neuron density and 100 μm for astrocyte reactivity ^42^. This is more damage than expected from most neuroscience tools such as a tetrode, silicon microelectrode, and especially fine, flexible single cell tools like the NET probe ^43^. For DISC the goal is to record LFPs, which are naturally more stable in chronic applications than single units ^44, 45^. Only the most local LFP sources should be affected. Future work should conduct long-term recordings but also explore the parameters of resistivity and thickness inside of a more detailed biophysics/FEM model (Suppl. Fig. 3) ^22^. Based on the work of others, recording of multi-unit cells in humans is also possible with even larger depth electrodes (1.2 mm Ø) typically 5 days after surgery (presumably after edema is cleared) ^46, 47^. The amplification of the signal due to the substrate insulation should increase the recording radius of even a small dipole moment from 125 μm to 195 μm (Suppl. Fig. 10). We do not suggest the use of DISC in mice given the prior lack of usefulness from other large devices such as a U-probe (300 μm diameter). The small twist-drill hole we use (0.9 mm) may have been an important factor for its success in rats. Since our experiments were acute, we used a stainless steel (SS316) core, although a silicone- or polyurethane cylinder would provide greater flexibility and that may result in greater signal stability in long-term applications ^48^. Future work will use a stiff guidewire for the initial insertion followed by a flexible version of DISC, made using advanced manufacturing methods and materials.

A particular advantage of DISC is its similar shape to the stereo-EEG depth array, enabling its use in clinical environments with little alteration. Relative to grid arrays, DISC may also enable more precise localization of seizures, and safety relative to grid arrays, as reviewed earlier. Thousands of epilepsy patients receive 8-20 sEEG devices each year for typically 1-2 weeks ^49^. In one study reviewing 150 procedures, the robotic sEEG surgery averaged 81 minutes and 6.2 minutes per electrode ^29^. The average surgery in this review implanted 13 sEEGs, having an insertion angle of 0-75° with no significant change in registration accuracy based on angle. By contrast, the Utah array has only been implanted in approximately 21 persons with 1-6 implants each ^50^. A single Utah array requires approximately 30-70-mm diameter skull opening for insertion. The cumulative neurosurgical experience and exceptionally low infection and hemorrhage rate when using sEEGs offer a safety advantage to non-stereotactic approaches ^10^. We provide a summary of recent studies and meta-analyses in Suppl. Table 4, which highlights the safety advantages that many sEEGs (typically 10) have relative to one or two ECoG devices.

## Outlook

This work demonstrates a highly effective, compact tool for improving amplitude, SNR, and decoding accuracy following whisker stimulation and supported by extensive modeling. More research is needed to demonstrate source localization, especially for sources beyond a few millimeters. Two or more DISC arrays in tissue may be highly effective in future work and would benefit from finite element software like Brainstorm in source localization ^51^. We will continue to improve the manufacturability of DISC and demonstrate its diagnostic capability in epilepsy models and advanced decoding tasks. We hope this work encourages the field to accelerate the trend toward smaller ring electrodes, directional leads, and ultimately toward high-density circular (DISC-like) arrays to explore the information capacity of LFPs “in stereo” over multiple spatial scales.

## Methods

### 1. Modeling and simulation

ANSYS Electronics Desktop 2021 R1 with the DC Conduction module was used throughout this work. A tissue block was modeled as 14 x 14 x 14 mm^3^, σ_tissue_ = 0.2 S/m, ε_tissue_= 80 ^52^. Lead substrate was 6-mm tall and a variable diameter, Ø_sh_, between 10 and 1400 microns. DISC diameter was by default 0.8 mm to emulate a standard sEEG device. 12 rows (200 µm vertical pitch) with 8 columns (every 45 degrees) were chosen to span the voltage field of interest without compromising simulation time. A variable electrode size was also used, Ø_e_, between 10 and 1000 microns. Due to the cylindrical geometry around an 800 µm device, when Ø_e_ is greater than 400 microns, a ring electrode is formed. Electrode diameter was 50 µm by default. Σ_shaft_ = 1e-10 S/m. ε_shaft_ = 2.7. Cortical monopoles and dipoles had identical properties as tissue.

Murakami and Okada demonstrated an invariant current dipole moment across animals of *Q* = 1-2 nAm/mm^2^, where *Q = I_d_ x d* ^27^. We chose *Q* = 2 nAm/mm^2^ and a dipole distance, *d*, of 1.2 mm center to center as the rat cortex is 2.1 mm. Therefore, a uniform current density of *I_d_* = 1.67 μA/mm^2^ was applied to the surface of all sources.

Unless otherwise stated, the dipole diameter was 200 μm. All microelectrodes and ring electrodes were platinum, σ_Pt_= 9.3e6 S/m. The ring electrode was 0.4 mm tall. Three contiguous surfaces of the tissue block outer surfaces were defined as 0V to emulate a distant reference. Figure 2 demonstrated 8 unique sources spread radially using an arrangement shown in Fig. 2A,B with details in Supplemental Table 2. That arrangement was intended to capture micro-, meso-, and macroscale sources interacting within a range of 5.1 mm in all directions. Sources 1, 2, and 8, and separately 3 and 4 had overlapping angular positions and therefore used the same electrode column for best amplitude. For each trial, one primary source was assigned a phase, =0π, and frequency, *f*=16.67 Hz, and the other 7 sources were assigned a random phase (0π to 2π) and frequency (40 Hz to 150 Hz). Electronic noise is added to each device according to its electrode area and empirical data with electrodes of that size. Ring electrodes (0.4 mm tall) had a noise value of 2.7 µV RMS. DISC had 96 electrodes with an independently assigned noise value of 4.3 μV RMS. Both noise values were randomly generated using a Gaussian distribution where the standard deviation was set to the RMS noise value. The waveform and SNR calculations were made as follows:

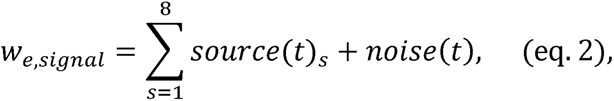

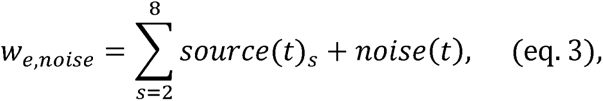

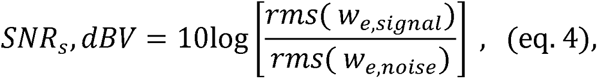

where index *e* is specific to one electrode and device, and index *s* is specific to a source. Source amplitudes were sampled at the surface of the electrode. Source 1 in equations 2, 3 is, by definition, the primary or phase zero source. The primary source rotates for each of eight positions in Fig. 2A. SNR was computed for one cycle of the primary source. The maximum amplitude of each source for each electrode and device type was computed by the ANSYS model as described before. The size and location of each source greatly influence the amplitude and noise contribution.

Suppl. Fig. 1 and Suppl. Movie 2 were a comparison of Ø_sh_ = 70 and 800 μm. Dipoles were either parallel or orthogonal and varied of a gap distance of 0.15 to 5 mm. The “large dipole” source was 800-μm diameter.

### 2. Fabrication and assembly

#### a. Microwires (Tetrode)

Tetrodes, each comprising eight insulated nichrome microwires (PX27, Sandvik AB, Sweden) were prepared following previously reported methods^53^. 35-40 turns on a vertical tetrode spinner (Neuralynx) ensured a compact tetrode of 8-cm length and approximately 50-μm diameter. Rigidity was enhanced by applying a temperature cured (250°F by a heat gun, 10 mins) thin layer of medical grade epoxy (Epoxy 301, Epoxy Technology), followed by connecting the tetrode to an Electrode Interface Board (EIB-36, Neuralynx). A nominal impedance (∼250 kΩ) was achieved by electroplating in a gold non-cyanide solution (Neuralynx) within an ultrasonic bath using an AutoLab Electrochemical Workstation (Metrohm).

#### b. 128-ch DISC assembly

Planar 128-channel arrays (G-128, Diagnostic Biochips, Inc) made of polyimide (HD Microsystems PI2611) formed the electrode body. G-128 model has 200 μm and 250 μm vertical and horizontal pitch, respectively, and follows a 16x8 matrix (row x column) configuration. Each electrode was 80 μm in diameter and was coated with PEDOT. The thickness of the polyimide substrate with electrodes was approximately 12 μm. The planar electrode array was wrapped onto a stainless-steel guidewire (GWXX-0170, 0.017” diameter, Component Supply Company) using several techniques (Fig. 7A-C) to create a cylindrical lead body. To eliminate the possibility of the guidewire interfering with the recording, PEBAX heat shrink tube (P2-023-0035-BLK, Cobalt Polymers) was applied on the SS wire and the tip was far from the recording sites. A 20-μm thick double coated acrylic tape (UTD-20B(W), Nitto) was utilized to wrap the flexible electrode array on the PEBAX coated guidewire. A FusionPro 48 Laser Machine (Epilog Corporation) was employed to pattern the thin adhesive film to match the shape of the G-128 probe. The thin adhesive film was placed on a flat substrate and fixed each corner with tape. Using the laser in vector mode at 60% speed, 6% of power, and 70% frequency with 2 cycles, the desired adhesive film area could be released within 20 seconds. Using a second heat shrink tube (PBST2-040-40-004C, Component Supply Company), the electrode array was pressed firmly to the guidewire to maintain better adhesion and proper geometry. The guidewire tip was carefully machined to create a smooth insertion end for implantation with minimal blood vessel damage. DISC pre-implantation impedance was found to be (235 ± 244kΩ), while post implantation impedance was (432 kΩ ± 1.84MΩ).

### 3. Animal surgical procedure

A total of 9 Sprague Dawley rats (250-450 g) were used in this study. Rats were housed in pairs in a regular 12 h/12 h light/dark cycle with ad libitum access to food and water. All experimental procedures, including analgesics and anesthesia, were approved by the Institutional Animal Care and Use Committee at Rice University. Rats were induced with 3-4% isoflurane (SomnoSuite, Kent Scientific) and all whiskers on the right facial pad were trimmed except B1, B2, C1, C2, D1, D2, E1, beta, delta, gamma, which were selected for stimulation. Subjects were mounted firmly with ear bars on a stereotactic frame (Digital U-Frame, Harvard Apparatus). Lubricating eye gel was used to keep the animal eyes hydrated. Topical lidocaine gel was applied on the ear bars to reduce pain during mounting and subsequent processes. Meloxicam (5mg/ml) with a dose of 2mg/kg was injected SQ as an analgesic. Animal body heat was maintained with isothermal heating pads. Hair was trimmed, and post application of topical ethanol and betadine, a 2-3cm rostro caudal incision was made to expose the skull. Hydrogen peroxide was applied to clean excess tissue and periosteum, while electrocautery was used to stop any unwanted bleeding. Anesthesia was maintained with 1-2% isoflurane during the surgery.

A Harvard Apparatus digital stereotactic frame with 3 axis readout was used to measure bregma and lambda coordinates for each subject. Using a micro burr (FST), a 4x4 mm^2^ area of the skull was thinned on the left hemisphere. Brain images through the cranial window were captured with a CMOS sensor camera (CS126MU, Thorlabs) mounted on a microscope (AmScope) using a 540 nm light source to add contrast to the subdural vessels. Major blood vessels were avoided, and a 900-um twist drill hole was made near our estimated coordinates relative to bregma. We assumed C1 was at -3.06 AP, 4.92 ML and D1 at -2.72 AP, 4.65 ML to help guide our selection. These positions were adjusted proportionally for the nominal Bregma-Lambda distance of 8.8 mm. A stainless-steel bone screw (#0-80, Grainger) attached with a 32 AWG copper wire served as a reference electrode implanted above the cerebellum. Tetrodes were implanted 1.4 mm deep, and DISC was implanted until the proximal-most electrode was in the brain (3.1-3.5 mm). Electrical isolation from EMG was achieved with a layer of dental acrylic at the bone ridge and silicone adhesive over the cranial burr hole (Kwik-sil; World Precision Instruments), and the reference electrode was isolated using dental acrylic.

### 4. Electrophysiology recording protocol

The whiskers were stimulated using a pneumatic dispenser (Nordson EFD Performus X). Each whisker was deflected by attaching a custom nozzle head to a standard 5mL syringe placed 2.5 mm from root of the whisker. The air puff (10 psi) from the nozzle head created a 10ms duration mechanical deflection in dorsal to ventral direction at 3 Hz over a 6s stimulus and 6s non-stimulus window; with a total of 450 stimulations recorded for each corresponding whisker. The whisker stimulation was driven by a battery-powered Arduino and MOSFET connected to our dispenser, with timing recorded by the Intan recording system. Wideband signals were amplified, digitized and recorded using a 256-channel Intan RHD interface board (Intan Technology) using a sampling rate of 20,000 Hz for both microwire and DISC electrodes. Recordings occurred in a Faraday cage unless otherwise noted. Neural signals were analyzed using offline custom Matlab scripts (Mathworks) and Python for figure generation.

### 5. Data processing

Raw neural signals were down sampled at 2 kHz, low-pass filtered (120 Hz, local field potential (LFP)), and band-pass filtered (40-150 Hz, gamma band). To account for signal variability, LFP and gamma signals were isolated for each trial. For each electrode, the LFP and gamma signal were averaged from across 450 trials. The signal amplitude was calculated by finding the distance between peak and valley poststimulation of the average signal. Maximum amplitude was found in DISC by looking at all electrodes used in infragranular layer. The best electrode was chosen as the maximum amplitude value from all whiskers. The SNR was computed using equation 4 with the rms signal post stimulus and the rms noise pre stimulus of the average signal.

Directional curves (polar plots) are defined by the peak-to-trough LFP amplitude and electrode column in one electrode row positioned at an infragranular layer. A unique column angle was noted for each device we assembled relative to the plane of the PCB. The plane of the PCB was always implanted parallel to the coronal plane. This angular information was used in the generation of polar plots. We matched the electrode amplitude of the average waveform to its angle. If an electrode was missing, the cubic interpolation was performed. A 16-point cubic interpolation was made from 8 electrode columns assuming equidistance between each column pair. Directional curves were standardized by subtracting the minimum amplitude.

Resultant vectors (RV) were calculated by the angle position of the electrode and the peak-to-peak LFP amplitude. The 8 original electrode positions were interpolated as described before to create 16 positions. The vectors were summed to create the resultant vector (length and angle). A ring electrode was simulated by averaging the signal from the electrodes in the DISC array. For a 2 mm virtual macroelectrode, we averaged all the channels that were included in a 2 mm vertical span (11 rows) of the DISC array, from the top of the brain to 2 mm deep in the brain. For the 0.4 mm virtual macroelectrode, we averaged all microelectrodes in 3 adjacent rows. The 0.4 mm rings were chosen to span supragranular, granular, and infragranular layers. The gap between each 0.4 mm ring was 0.4 mm.

We extracted the gamma band waveform from 0 to 100 ms poststimulation for each trial. The trial waveform from 44 electrodes, corresponding to 11 rows and 4 columns, were concatenated into one matrix and the mean subtracted. This was defined as our feature matrix with columns as the gamma waveforms, and rows as individual trials. To reduce the dimensionality of our feature matrix, Principal Component Analysis (PCA) was performed, choosing the number of components necessary to represent the 99% variance of our data. The reduced feature matrix was then divided into 10-folds. Each fold used 80% for training and 20% for testing with a linear discriminant analysis (LDA) model.

By calculating the second derivative of the average LFP waveform, we computed the current source density (CSD, ^54^) of each DISC row. For optimal results, the 20,000 Hz sampling rate waveform was used.

### 6. Statistical analysis

Generalized Linear Mixed Model (GLMM) was used to evaluate the differences of DISC device (intercept) compared to tetrode and virtual rings. The subject ID was used as a random effect, and SNR, amplitude, or model accuracy as dependent variables. DISC was the defined baseline in the GLMM. When comparing between any two groups besides DISC, Tukey’s range test was employed.

### 7. Perfusion and Histology

A subset of rats had histology performed to identify the barrel location. Rats were euthanized by applying 5% isoflurane as a primary method, followed by cervical dislocation and thoracotomy as a secondary method and confirmation, respectively. The subset prepared for histology were euthanized using 5% isoflurane and cervical dislocation, followed by transcardial perfusion of 2% PFA.

The cortex was flattened by scooping out the thalamus and placing in between two glass slides ∼1.5mm apart. The flattened brain was kept in 1x PBS for 6-8 hours at 4°C. It was later transferred to a 2% PFA solution for 24 hours. Fixed flattened brain was washed with 0.1M 1X PBS and was sectioned using a vibratome (Leica VT 1200S, Germany) to create 50-80-μm slices. Sections were transferred to a culture plate culture plate and rinsed in HEPES buffer (0.1M pH 7.4) for 15 minutes. Cytochrome oxidase staining solution was prepared similar to [^55^] and the sections were incubated in the solution at 37° C. Stains were visible around 30-60 mins. 2% PFA was added with the solution to stop the staining reaction. Sections were mounted on microscope glass slides and rinsed in consecutive 70% (2 mins), 96% (2 mins) and 99.5% (3 mins) ethanol. Sections were rinsed in isopropyl alcohol for 3 mins, followed by xylene rinse for 5 minutes to complete the dehydration process. Applying a non-aqueous mounting media, the sections were cover-slipped and imaged using a microscope (Keyence BZ-X800, Japan) and 20X objective. Electrode implantation location was reconstructed from the slides containing stained barrel cortex maps.

## Data Availability

All datasets generated and analyzed during the current study will be made available upon publication.

## Code Availability

Custom Python and Matlab code for data analysis and the instructions for use can be found at https://github.com/TBBL-UTHealth/DISC1.

## Acknowledgements

We would like to acknowledge John Mosher (UTHealth) for helpful discussions on source localization and dipole modeling. We thank the staff at Rice University’s nanofabrication facility for supporting our neural probe development. We thank Mihály Vöröslakos for sharing helpful surgery tips and Charles Schroeder for his insights into local field potential recording with varied devices. We acknowledge the Texas Institute of Restorative Neurotechnologies at UTHealth for its financial support.

## Authorship Contributions

J.P.S. and N.T. conceived of the presented idea. A.M.A. led the analytical methods and machine learning, and M.R.I. and J.P.S. verified results. W.K. led the surgeries, histology, and device assembly and X.B. assisted with device assembly. C.W. led the finite element modeling and J.P.S. assisted. M.R.I. led the single statistic classification modeling. M.G. led the fabrication and mechanical modeling. All authors discussed the results and contributed to the final manuscript. J.P.S. supervised the project.

## Extended Data

**Extended Data Table 1.** LDA Classification Accuracy by Device and Electrode Configuration. **Note a:** For the 11xn analysis, we selected 11 contiguous rows in the cortex, generally rows 1-11 or 2-12. Generally, the deepest rows showed little amplitude or variation. 3xn analysis selected 3 depths approximately inside layers II/III, V, and VI using depths from Paxinos, Watson 6^th^ Edition, Figure 52. 1xn analysis used layer V recordings. Each subject was adjusted slightly based on row 1 impedance and the location of a polarity change. **Note b:** Best channel was chosen as the channel in layer V with the highest amplitude as verified with a negative peak. This electrode channel was fixed for each subject with no change by whisker. **Note c:** Accuracy after 10-fold cross-validation.

| Device | Electrode Configuration (Rows x columns) | Subject Accuracy (%)<sup>c</sup> |  |  |  |  |  |  |  |  | Average Accuracy (%) |
| --- | --- | --- | --- | --- | --- | --- | --- | --- | --- | --- | --- |
|  |  | S1 | S2 | S3 | S4 | S5 | S6 | S7 | S8 | S9 |  |
| **DISC<sup>a</sup>** | 16x8 | 90.2 | 98.6 | 99.1 | 95.9 | 94.0 | 93.2 | 91.5 | 99.2 | 95.9 | 95.3 |
|  | 16x4 | 92.5 | 98.2 | 98.2 | 95.4 | 91.3 | 90.9 | 89.9 | 97.7 | 94.7 | 94.3 |
|  | 11x8 | 92.7 | 98.4 | 98.5 | 96.3 | 92.3 | 93.5 | 91.6 | 98.8 | 95.7 | 95.3 |
|  | 11x4 | 93.8 | 97.7 | 92.3 | 95.5 | 89.3 | 91.4 | 89.8 | 97.7 | 94.7 | 93.6 |
|  | 11x1 | 38.3 | 65.8 | 63.4 | 90.2 | 66.8 | 84.7 | 54.0 | 84.6 | 71.3 | 68.8 |
|  | 3x8 | 92.8 | 97.1 | 84.9 | 93.8 | 88.9 | 88.7 | 86.5 | 97.8 | 92.0 | 91.4 |
|  | 3x4 | 92.7 | 94.5 | 81.5 | 91.8 | 87.3 | 85.9 | 83.6 | 95.4 | 91.1 | 89.3 |
|  | 1x8 | 57.8 | 82.0 | 61.9 | 85.9 | 72.1 | 79.4 | 82.6 | 96.1 | 85.3 | 78.1 |
|  | 1x4 | 57.3 | 57.1 | 48.0 | 85.0 | 68.0 | 75.6 | 77.9 | 91.4 | 80.8 | 71.2 |
|  | Best 1x1<sup>b</sup> | 32.7 | 30.6 | 38.2 | 64.3 | 42.6 | 59.3 | 30.4 | 56.6 | 50.2 | 45.0 |
| **Tetrodes** | Best 4 | 48.7 | 50.5 | 72.6 | 57.2 | 26.7 | 30.3 | 51.0 | 89.3 | 95.3 | 58.0 |
|  | Best 1 | 38.6 | 44.0 | 29.3 | 50.4 | 21.2 | 28.7 | 39.5 | 60.2 | 86.1 | 44.2 |
| **Virtual Macro** | 2 mm | 25.1 | 41.5 | 37.5 | 73.8 | 22.4 | 41.0 | 25.1 | 34.7 | 31.6 | 37.0 |
|  | 3 x 400 um | 35.8 | 63.7 | 57.4 | 81.6 | 51.1 | 63.5 | 65.6 | 69.6 | 54.4 | 60.3 |

**Extended Data Fig. 1:**
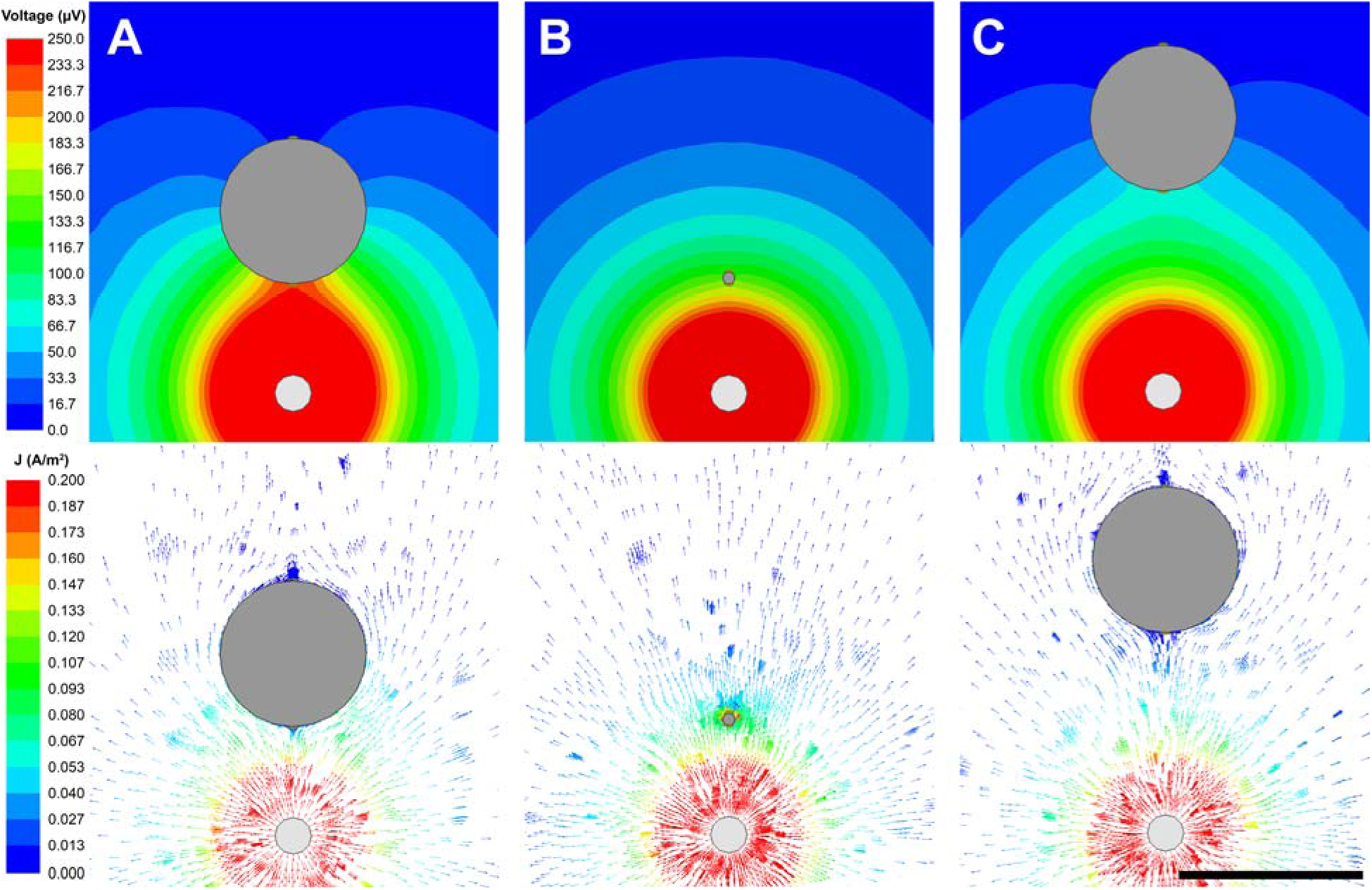
2D Voltage contour and J-field illustrating the mechanism of substrate shielding. Top-down view of a current field (bottom) and potential (top) contours around insulating bodies. Scale bar: 1 mm. **A,** A dipole and 800-μm diameter lead body (gray) with a 500 µm gap. **B**, 65-μm lead body (microwire-sized device) with a 500 µm gap. **C,** 800-μm diameter lead body with 1-mm gap. The relative positions and scales are kept constant in rows 1 and 2. The current vectors (right) illustrate the rapid divergence around the insulating body compared to microwire-sized array (row 2). The potential contour lines (left) are orthogonal to the J-field. The voltage gradient (density of contour lines) is unchanged where the change is current is near zero. Thus, an isopotential extends away from the electrode toward the source in response to a static J-field. Similarly, an isopotential extends away from the opposite side electrode and has a much lower magnitude. In summary, DISC shows greater directionality (potential differences) as the device diameter increases, directing more current flow around the lead body shaft and amplifying the difference in front and back electrode potential.

**Extended Data Fig. 2.**
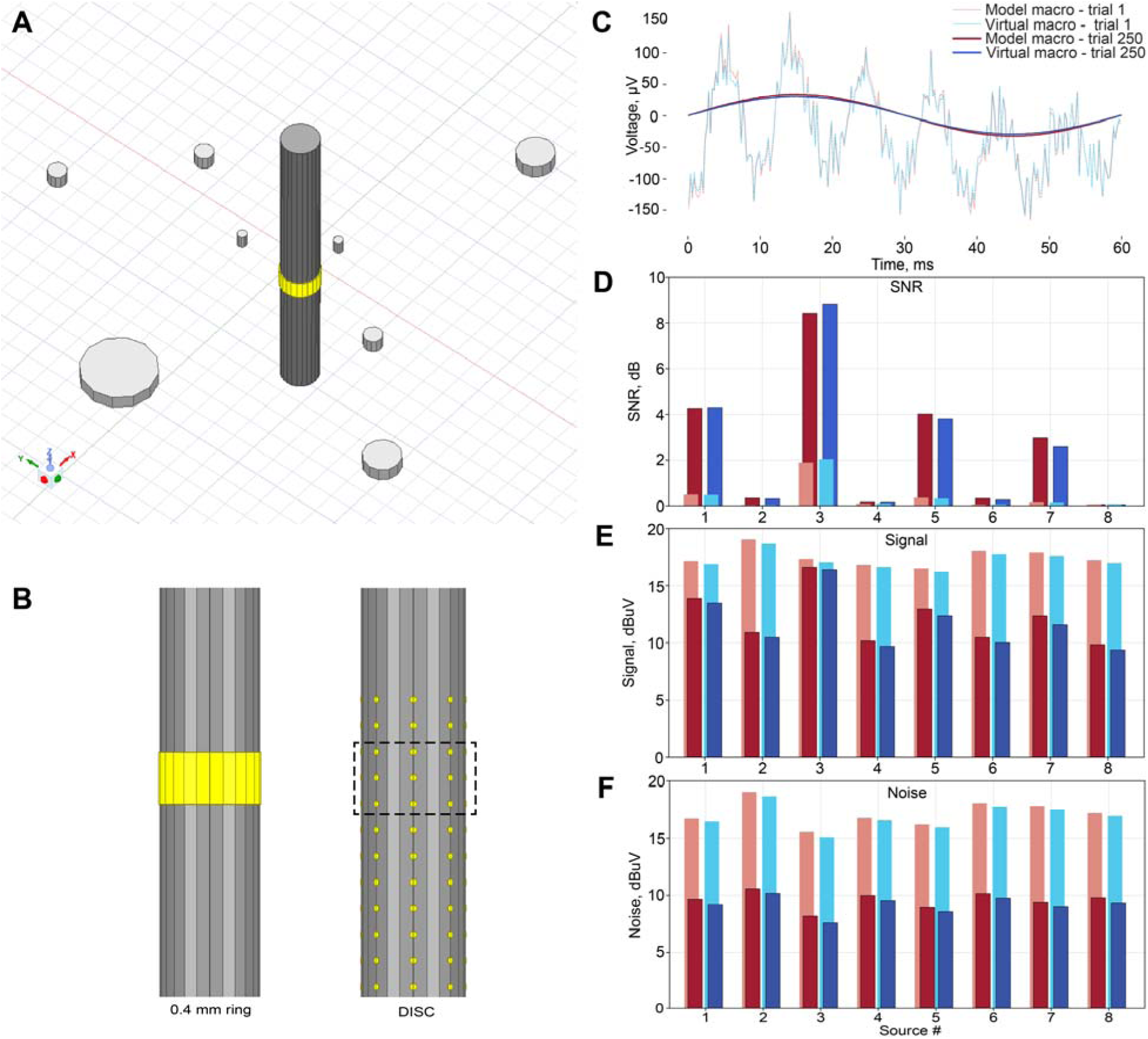
SNR, amplitude, and noise comparison between a FEM ring and virtual DISC ring electrode. **A,** FEM model of a 0.4-mm tall ring electrode surrounded by 8 unique dipoles (only the sink of dipole is shown), **B,** and a virtual 0.4-mm ring formed from a 3 x 8 microelectrode array (middle) are juxtaposed in an 8-source environment. Both devices were modeled using Suppl. Table 1, as was used in Fig. 2. **C,** Waveforms for the single trial and averaged trials. **D,** The SNR, **E,** amplitude, and **F,** noise are separated for comparison. The left column represents one randomly seeded trial, and the right is the average after 250 trials (saturated color). The ring was assigned 2.7 μVrms while each microelectrode was assigned 4.3 μVrms as before, which represented the average noise floor values. SNR, RMS signal and RMS noise were calculated using equations 4, 2, and 3, respectively. These results indicate a deviation of no more than 0.5 dB or dBV for any measurement. Source phase, frequency, and noise were kept consistent between models for trial 1, but the remaining 249 trials were randomized.

**Extended Data Fig. 3.**
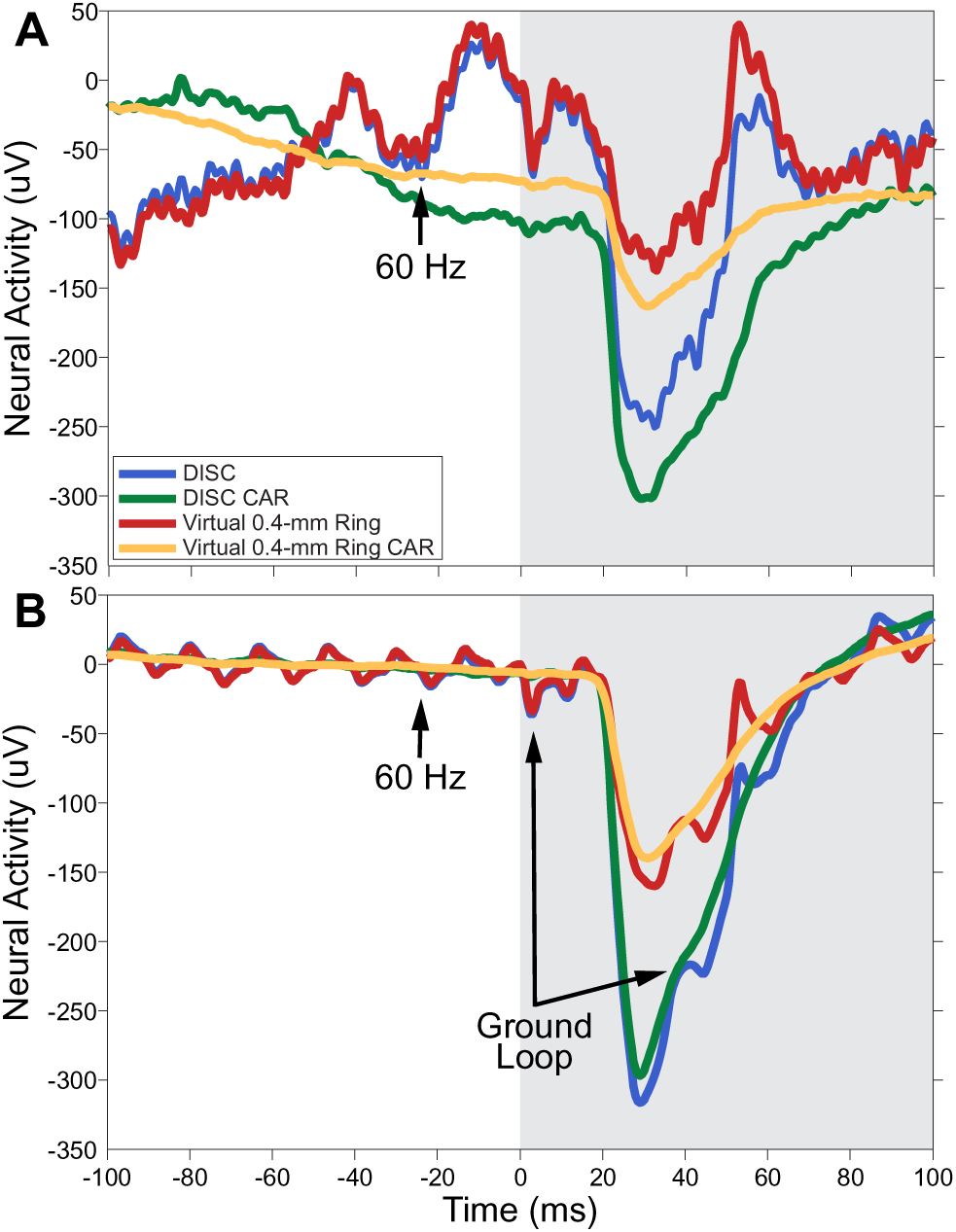
Noise immunity of microelectrodes using common average referencing (CAR). **A,** Sample single trial of DISC in an *in vivo* stimulation trial without the use of a Faraday cage. The ground loop was intentionally created by connecting the air puffer ground at a separate outlet. All signals were identically filtered. CAR results in a small decrease in the evoked potential but removes the visible common mode noise. A virtual ring electrode was computed from a 3 x 8 configuration and referenced to the average of all remaining 104 electrodes.

## Supplementary Information

### Contents

**Suppl. Table 1a.** Source configuration table for a multi-source model (see Fig. 2).

| Source number | Diameter, $\varnothing_{\text{source, } \mu\text{m}}$ | Gap distance, mm | Angle, degrees |
| --- | --- | --- | --- |
| 1 | 200 | 1 | 0 |
| 2 | 400 | 3.16 | 19 |
| 3 | 100 | 0.6 | 94.4 |
| 4 | 800 | 5.12 | 107 |
| 5 | 400 | 1.73 | 190 |
| 6 | 800 | 4.63 | 212 |
| 7 | 1600 | 3.93 | 345 |
| 8 | 400 | 5.0 | 349 |

**Suppl. Table 1b.** Electrical properties of electrostatic model (see Fig. 2).

| Lead Height ($\mu\text{m}$) | Lead Conductivity (S/m) | Brain Conductivity (S/m) | Electrode Conductivity (S/m) | Electrode Pitch ($\mu\text{m}$) | Electrode Radius ($\mu\text{m}$) | Current Density, $\mu\text{A}/\text{mm}^2$ |
| --- | --- | --- | --- | --- | --- | --- |
| 6000 | 0.1e-9 | 0.2 | 9.3e6 (platinum) | 200 | 25 (default) | 1.67 |

**Suppl. Table 2.** Best Electrode SNR, Best Electrode Signal RMS, and Best Electrode Noise RMS of LFP band by Device.

| Device | Electrode Configuration (Rows x columns) | SNR LFP (dB) |  |  |  |  |  |  |  |  | Average SNR |
| --- | --- | --- | --- | --- | --- | --- | --- | --- | --- | --- | --- |
|  |  | S1 | S2 | S3 | S4 | S5 | S6 | S7 | S8 | S9 |  |
| **DISC** | 11x4 | 10.8 | 16.8 | 14.6 | 13.2 | 10.0 | 12.3 | 10.3 | 14.2 | 14.8 | 12.6 |
| **DISC CAR** | 11x4 | 13.9 | 18.3 | 15.9 | 14.5 | 13.7 | 15.3 | 15.5 | 15.2 | 15.5 | 15.3 |
| **Tetrodes** | Best 4 | 8.0 | 12.3 | 7.5 | 10.4 | 5.8 | 5.1 | 6.8 | 7.6 | 7.3 | 7.7 |
| **Virtual Macro** | 2 mm | 3.8 | 12.2 | 8.8 | 11.1 | 4.0 | 4.9 | 2.8 | 8.5 | 9.8 | 7.1 |
|  | 3 x 400 um | 8.1 | 14.1 | 12.7 | 11.8 | 6.6 | 9.5 | 7.0 | 11.8 | 11.5 | 10.1 |

| Device | Electrode Configuration<br>(Rows x columns) | RMS Signal (uVrms) |  |  |  |  |  |  |  |  | Average RMS Signal |
| --- | --- | --- | --- | --- | --- | --- | --- | --- | --- | --- | --- |
|  |  | S1 | S2 | S3 | S4 | S5 | S6 | S7 | S8 | S9 |  |
| **DISC** | 11x4 | 20.7 | 73.1 | 46.8 | 73.1 | 10.9 | 25.4 | 12.3 | 29.0 | 41.4 | 35.2 |
| **DISC CAR** | 11x4 | 18.8 | 61.3 | 42.4 | 32.4 | 9.4 | 22.3 | 11.3 | 24.8 | 33.5 | 27.8 |
| **Tetrodes** | Best 4 | 24.1 | 56.3 | 28.9 | 18.7 | 10.1 | 19.5 | 20.5 | 28.7 | 42.1 | 25.3 |
| **Virtual Macro** | 2 mm | 9.3 | 41.3 | 18.5 | 67.9 | 6.1 | 11.7 | 4.2 | 13.8 | 28.7 | 21.5 |
|  | 3 x 400 um | 24.0 | 93.0 | 69.3 | 93.6 | 13.1 | 36.8 | 11.7 | 37.3 | 50.8 | 44.6 |

| Device | Electrode Configuration<br>(Rows x columns) | RMS Noise (uVrms) |  |  |  |  |  |  |  |  | Average RMS Noise |
| --- | --- | --- | --- | --- | --- | --- | --- | --- | --- | --- | --- |
|  |  | S1 | S2 | S3 | S4 | S5 | S6 | S7 | S8 | S9 |  |
| **DISC** | 11x4 | 4.0 | 4.5 | 4.5 | 7.2 | 2.4 | 4.9 | 2.7 | 3.1 | 4.5 | 4.1 |
| **DISC CAR** | 11x4 | 2.4 | 3.6 | 3.7 | 3.5 | 1.1 | 2.6 | 1.1 | 2.6 | 2.9 | 2.6 |
| **Tetrodes** | Best 4 | 4.6 | 4.5 | 5.9 | 2.5 | 3.4 | 6.0 | 4.3 | 5.4 | 9.2 | 5.4 |
| **Virtual Macro** | 2 mm | 3.3 | 2.6 | 2.5 | 6.0 | 2.1 | 4.1 | 2.5 | 2.0 | 3.4 | 3.1 |
|  | 3 x 400 um | 3.8 | 4.3 | 4.3 | 6.8 | 2.4 | 4.8 | 2.7 | 3.0 | 3.9 | 3.9 |

**Suppl. Table 3.** LDA classification accuracy (%) with different types of individual statistics estimated from gamma-band, DISC and tetrode. *Note on Table 3:* In another effort to study the contributing factors in decoding, a simplified feature matrix was created using the following scalars: Zero-crossing, RMS, variance, Phase at maximum peak of Fourier transform (FT), resultant vector, Power at maximum frequency spectrum (PMFS), Frequency (Freq) at maximum peak, and Time Delay (TD). We used 11 x 4 configuration in this analysis using gamma band response (0:0.5:100 ms). We performed 10-fold cross-validation with LDA such that each run serves once for validation. Finally, the classification result with accuracy was taken by averaging all the runs.

| Device | Method | Subject-wise Accuracy (%) |  |  |  |  |  |  |  |  | Avg Accuracy (%) |
| --- | --- | --- | --- | --- | --- | --- | --- | --- | --- | --- | --- |
|  |  | S1 | S2 | S3 | S4 | S5 | S6 | S7 | S8 | S9 |  |
| **DISC** | ZC | 20.1 | 13.7 | 11.3 | 32.6 | 9.14 | 13.0 | 35.6 | 27.7 | 13.1 | 19.6 |
|  | **RMS** | **43.0** | **83.0** | **82.5** | **70.5** | **67.1** | **56.4** | **53.9** | **41.6** | **67.4** | **62.8** |
|  | Var | 41.4 | 56.5 | 72.3 | 66.7 | 56.5 | 35.7 | 44.6 | 31.7 | 56.5 | 51.3 |
|  | Phase | 12.6 | 61.5 | 32.4 | 27.1 | 35.6 | 43.3 | 16.6 | 38.9 | 19.9 | 31.9 |
|  | RV | 31.9 | 32.8 | 38.9 | 42.3 | 33.7 | 24.7 | 34.4 | 22.9 | 53.1 | 34.9 |
|  | PMFS | 42.1 | 72.3 | 58.2 | 69.4 | 59.1 | 63.0 | 44.5 | 40.3 | 57.3 | 56.2 |
|  | Freq | 29.7 | 28.8 | 27.0 | 23.4 | 26.6 | 14.0 | 29.9 | 19.6 | 5.93 | 22.7 |
|  | TD | 13.3 | 40.7 | 30.6 | 34.7 | 40.5 | 39.3 | 15.43 | 22.0 | 25.9 | 29.1 |
| **Tetrode** | ZC | 3.2 | 3.3 | 17.9 | 11.6 | 12.5 | 3.0 | 11.7 | 3.8 | 12.3 | 8.80 |
|  | **RMS** | **27.9** | **32.5** | **14.4** | **32.5** | **28.6** | **12.7** | **15.0** | **22.9** | **26.5** | **23.6** |
|  | Var | 27.1 | 31.6 | 20.3 | 22.6 | 11.6 | 12.5 | 14.3 | 21.8 | 17.2 | 19.8 |
|  | Phase | 25.1 | 37.9 | 11.6 | 4.27 | 13.6 | 12.2 | 19.3 | 29.6 | 13.2 | 18.5 |
|  | PMFS | 28.5 | 31.2 | 20.3 | 31.4 | 22.9 | 20.3 | 22.8 | 13.2 | 17.9 | 23.2 |
|  | Freq | 25.3 | 20.9 | 10.5 | 20.0 | 4.54 | 11.4 | 20.9 | 21.1 | 22.4 | 17.4 |
|  | TD | 2.55 | 21.5 | 10.3 | 20.5 | 12.6 | 19.7 | 27.9 | 12.9 | 22.5 | 16.7 |

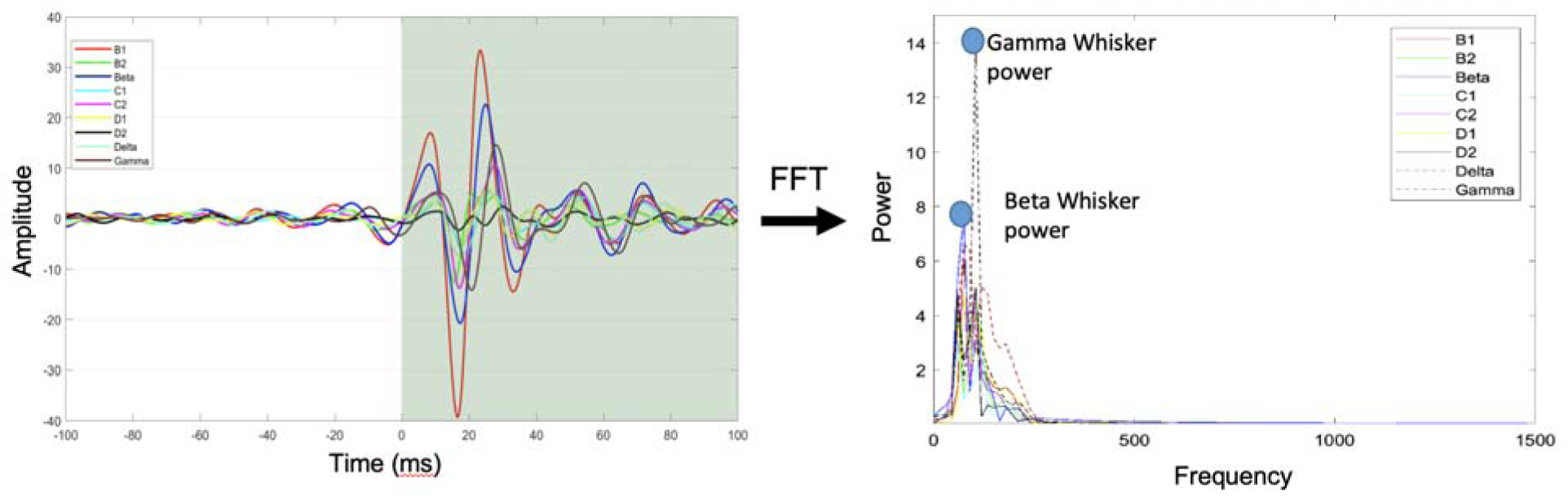

The short review of statistical methods is the following:

**Zero-Crossing (ZC):** The zero-crossing (ZC) is the number of sign changes along a Gamma LFP signal, defined as:

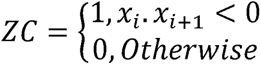

where *x_i_* and *x_i_*_+1_ are the two consecutive data points from a Gamma LFP signal.

**Root-Mean Square (RMS):** The root mean square (RMS) is amplitude from gamma LFP signals over 0-100 ms period:

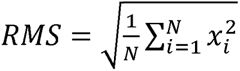

where *N* denotes the length of the signals *x* for each gamma band waveform

**Variance (Var):** The variance (Var) of signal refers to a statistical measuring how the amplitudes of the LFP signals for each whisker are spread out from their average value, and is also used to characterize the nature of each whisker LFP signals defined as:

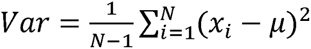

where *N* and μ represent the number of sample points and mean respectively from gamma band signal *x*.

**Phase:** In signal processing applications, the phase of signals may be another key feature for identifying whisker activity, here defined as:

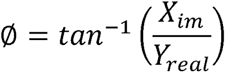

where *X_im_* and *X_real_* are the real and imagery parts of Fourier Transform (FT) at a maximum energy.

**Resultant Vector (RV):** For machine-learning application with DISC device, a set of vectors (rms and direction) was estimated from 8 electrode columns at layer V for each trial a to represent the resultant vectors. A simplified feature matrix is created based on the estimated resultant vector. An example of a resultant vector can be seen in Suppl. Fig. 8.

**Power at maximum frequency spectrum (PMFS)**: Absolute power at maximum peak from frequency spectrum of each trial was estimated using FT. Y-intercept on an FFT plot.

**Frequency (Freq):** Frequency at maximum power of each trial was estimated based on FT. X-intercept on an FFT plot.

**Time Delay (TD):** Time delay in a time domain was computed by taking distance between the onset of stimulus and highest gamma peak.

**Suppl. Table 4.**
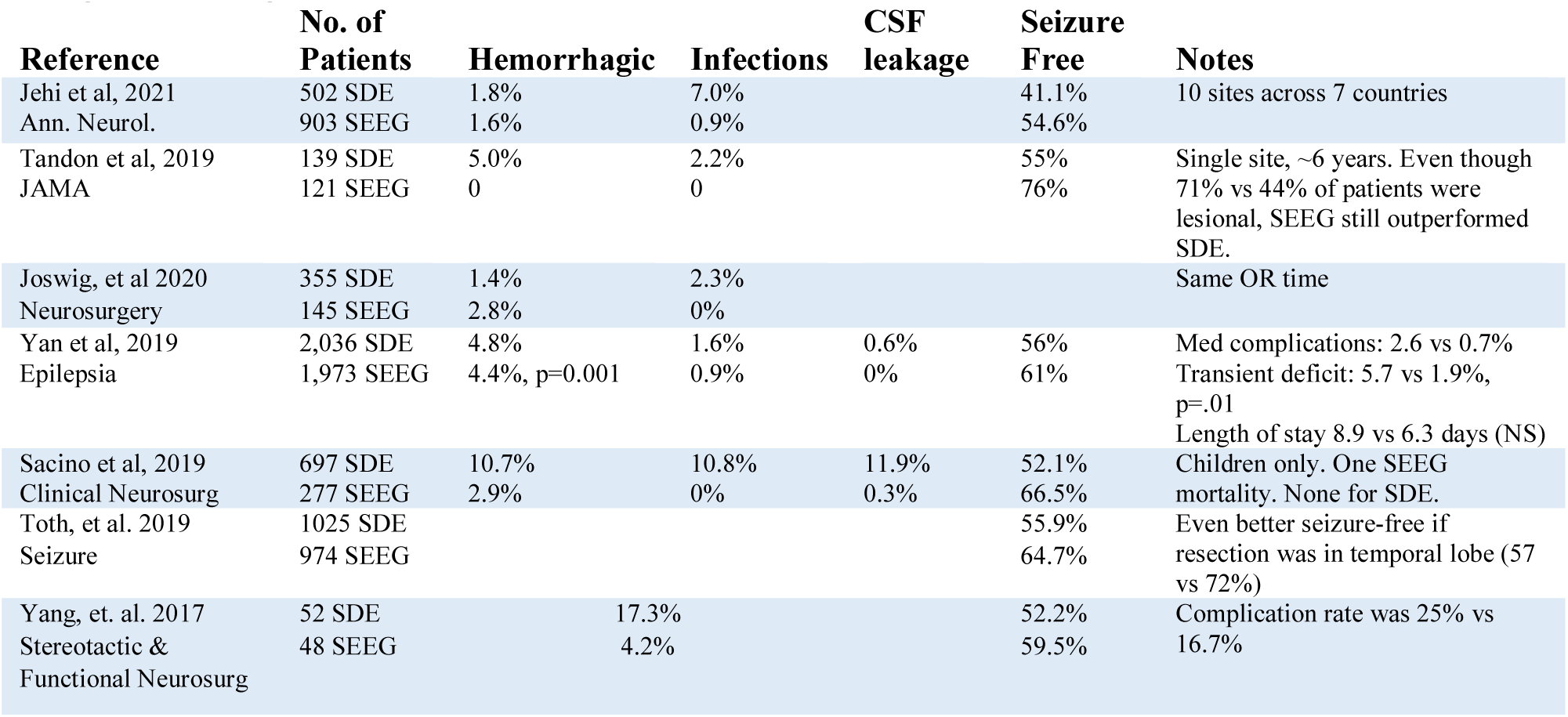
Review of Recent Safety and Efficacy Differences in Epileptogenic Diagnosis using ECoG vs SEEG. *Note on Suppl. Table 4*. Suppl. Table 4 lists 7 retrospective reviews comparing sEEG to ECoG implants. A reduced risk of infection was found in 6 of 6 studies and a reduction of hemorrhaging in 5 of 6. SEEG allows surgeons to reduce infection from 4.1 to 0.8% and hemorrhaging from 5.2 to 3.3% (weighted average by population size), which does not include the Yang 2017 study. sEEG surgeries typically use 8 to 15 devices to achieve high channel count and spatial diversity.

**Suppl. Fig. 1.**
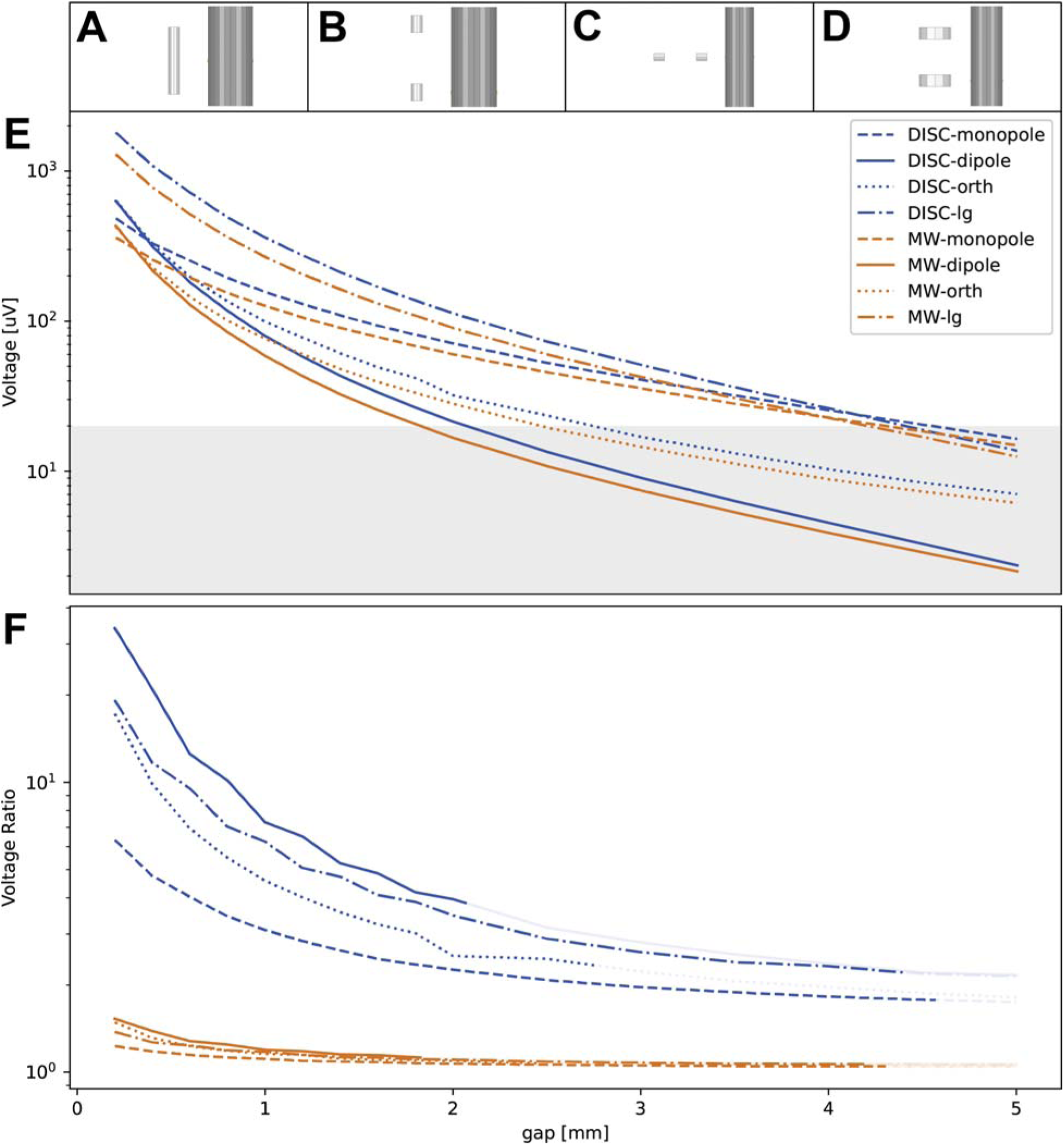
Absolute and relative voltage by source type and orientation for DISC and microwire-equivalent substrate (MW) (related to Suppl. Movie 2)**. A,** Monopole and DISC model orientation, **B,** parallel dipole. **C,** orthogonal dipole, and **D,** large (800 µm), parallel dipole. **E,** Voltage measured at front electrode with increasing gap. Below 20 μV, those sources will be difficult to separate from noise. **F,** Front/back electrode ratio with increasing gap. Lines become faint after falling below 20 μV. *Note on Suppl. Fig. 1*: Both the 65-μm and 800-μm Ø structures had electrodes distributed over the surface and the best electrode amplitude is plotted above. The monopole is plotted for reference to approximate 1/r falloff. The parallel-oriented dipole has the greatest directionality but the least range (200 and 800-μm Ø have an estimated range of 2.1, and 4.3 mm). The orthogonal dipole has less dynamic change across the 800-μm array, but the signal is approximately detectable at 2.8 mm relative to 2.1 mm for same dipole when parallel. The microwire-equivalent substrate was 65-μm diameter and had lower amplitude and directionality. Interestingly, the larger dipole has slightly less falloff (rate of attenuation vs distance) than the smaller source and maintains a high F/B ratio. Future modeling will examine the tradeoff in orientation and range, with the loss in dynamic voltage values when conducting source separation. (See also Suppl. Movie 2).

**Suppl. Fig. 2.**
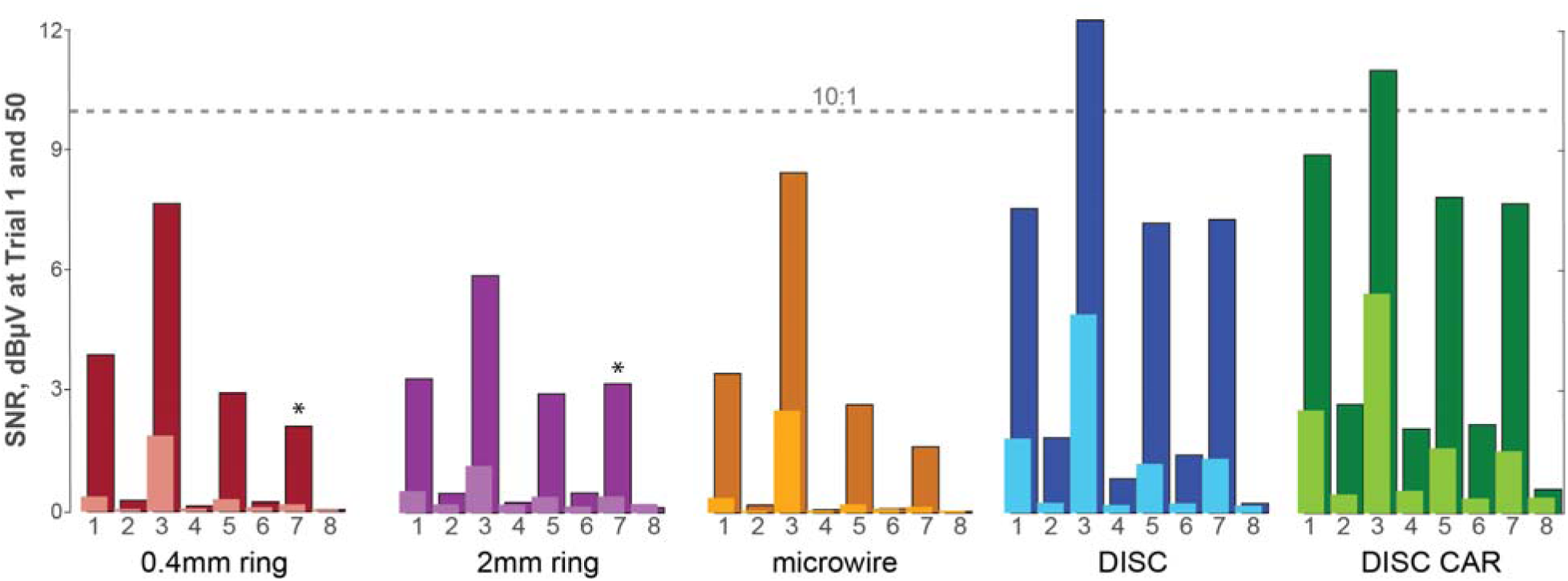
SNR results for an 8-source model with a 2-mm ring electrode. The FEM model described in Fig. 2 was also analyzed with a 2-mm tall ring electrode (standard size of an SEEG). Relative to the 0.4-mm-tall ring, this ring SNR was 1-2 dBµV or 25-50% lower in the typical case, and in the case of source 7 (large, distant neural source), the 2mm ring performs slightly better. This is because biological noise is attenuated much more for the larger ring, leading to better SNR values in the presence of high amplitude noise. All electrodes were placed in the same plane as the sink which provides a large amplitude for all electrode types. **(*)** The higher SNR value for sources 6 and 7 raises questions about the probability and scenarios when tall rings would be expected to outperform short ring electrodes (mini-sEEGs), which will be studied in future models.

**Suppl. Fig. 3.**
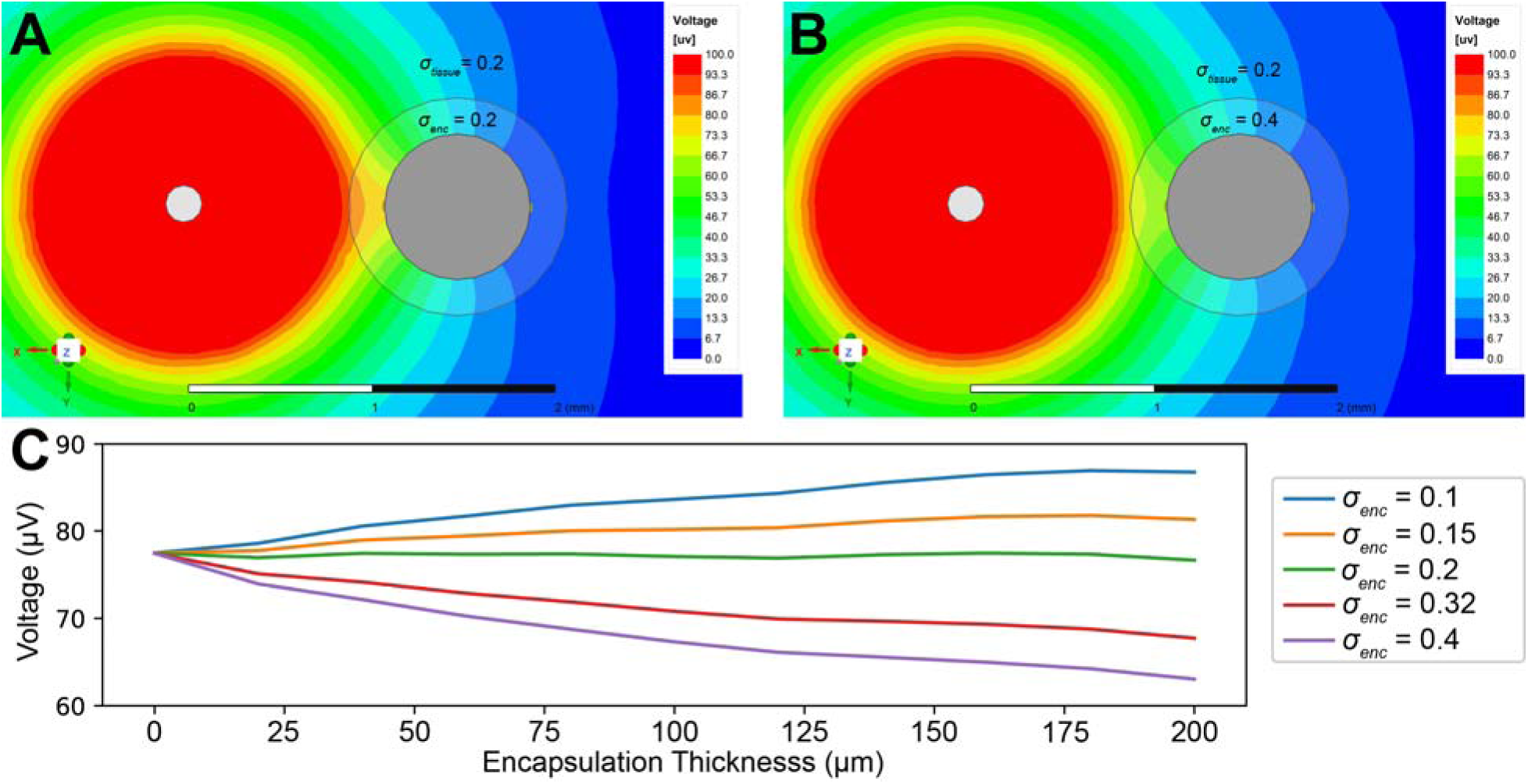
FEM model of encapsulation and edema around 800-μm diameter lead body. **A,** Voltage contours with a 0.2 S/m encapsulation layer, matching the brain conductivity. **B,** Voltage contours with a 0.4 S/m encapsulation layer, indicating edema formed around DISC. *Note on Suppl*. Fig. 3: Encapsulation thickness around the lead body was 0-200 µm thick and σ_enc_ 0.1, 0.15, 0.2, 0.32, and 0.4 S/m. A gap of 1 mm was used with a 200-µm diameter source. Even with a 0.4 S/m edemic layer, the amplitude reduction is less than 20%. While seemingly contradictory, encapsulation tissue amplifies the measured voltage as predicted originally by Moffitt, McIntyre 2005 ^1^. As illustrated in Extended Data Fig. 1, the voltage gradient (density of contour lines) is unchanged where the change is current is near zero. Encapsulation tissue in this range and at this thickness creates an extended isopotential in front of the electrode in response to a static J-field. The model used for this analysis is identical to the previous, with the addition of a cylindrical conductive layer around the device with the electrodes.

**Suppl. Fig. 4.**
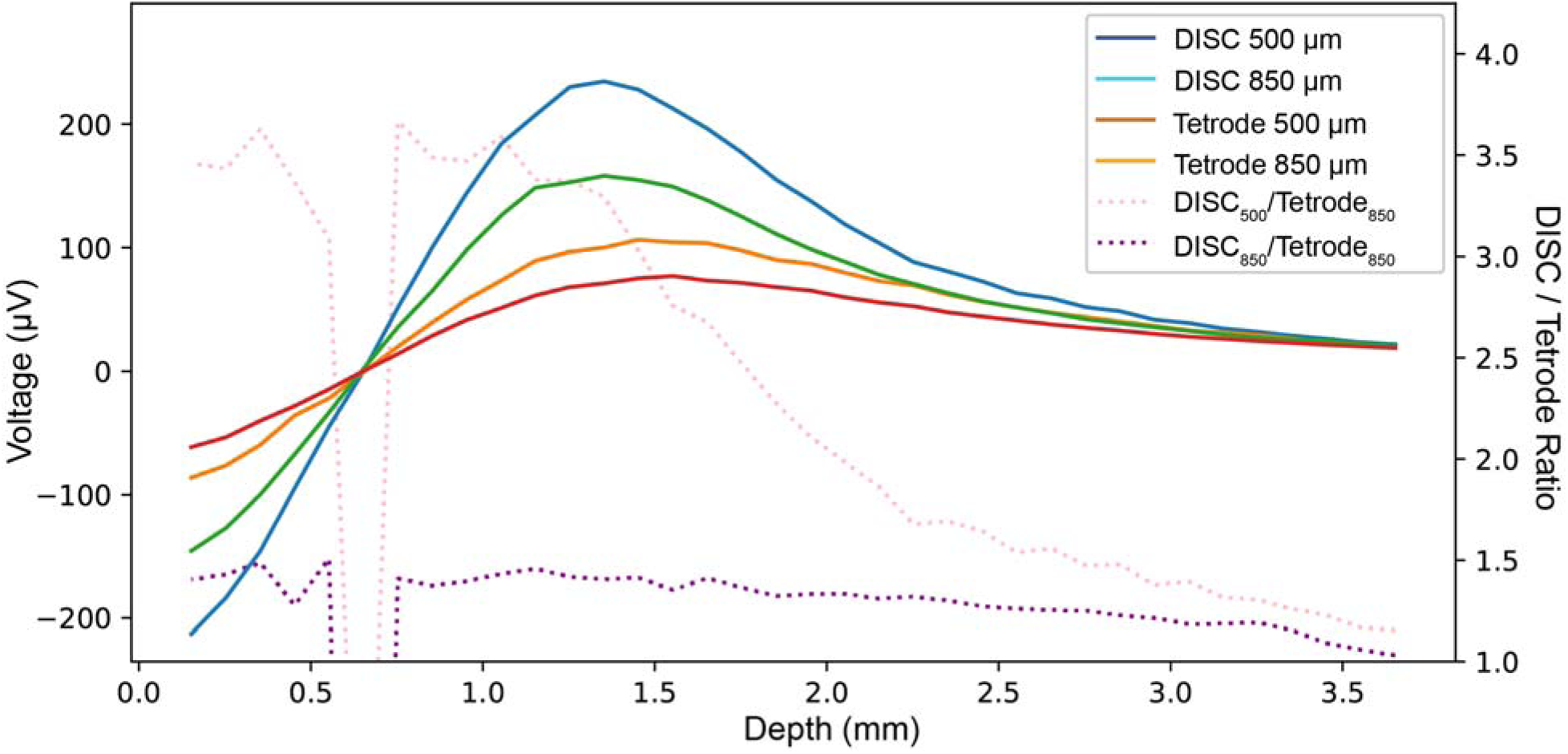
Voltage sensitivity analysis of depth in a recording array. FEM results of the absolute voltage measured along a parallel path of a current dipole source (200-μm Ø dipole source, 1.2 mm center-center sink to source as before, and a 500 or 850 µm gap) for both DISC (Ø: 800 μm) and a narrow substrate (Ø: 65 μm). The two ratios represent the approximate theoretical limits of amplification depending on how much closer DISC was to the source. *Note on Suppl.* Fig. 4: Our FEM model predicts an increase in amplitude of 41% (also shown in Fig. 1C) assuming a uniform gap between each electrode type and the source. However, the measured increase in amplitude was higher—3.14 ratio or 214% higher when comparing DISC (best infragranular electrode) to the tetrode (Fig. 3, using relative median values). *In vivo*, DISC is an unspecified distance closer to the surrounding cortical columns depending on the degree and gradient of compression after insertion, up to a maximum of 350 μm (DISC radius of 325-375 μm – radius of tetrode, 25μm). If our FEM model of DISC replaces the narrow substrate (microwire) and the sources do not move, then DISC will be 350 μm closer and measure a 217% increase. Unlike DISC, implanting the tetrode did not result in any noticeable dimpling so we believe the static depth of 1.4 mm used with tetrode measurements represents a value close to the maximum and was not highly sensitive to a slight shift in depth. Given the unknowns, we can only claim that a significant amplification was predicted, and the observed amplification approached the theoretical limit. Future studies with a Michigan-style array would add some precision to this amplification factor, but it would not be able to resolve the questions around lateral tissue compression (i.e. true distance to the sources).

**Suppl. Fig. 5.**
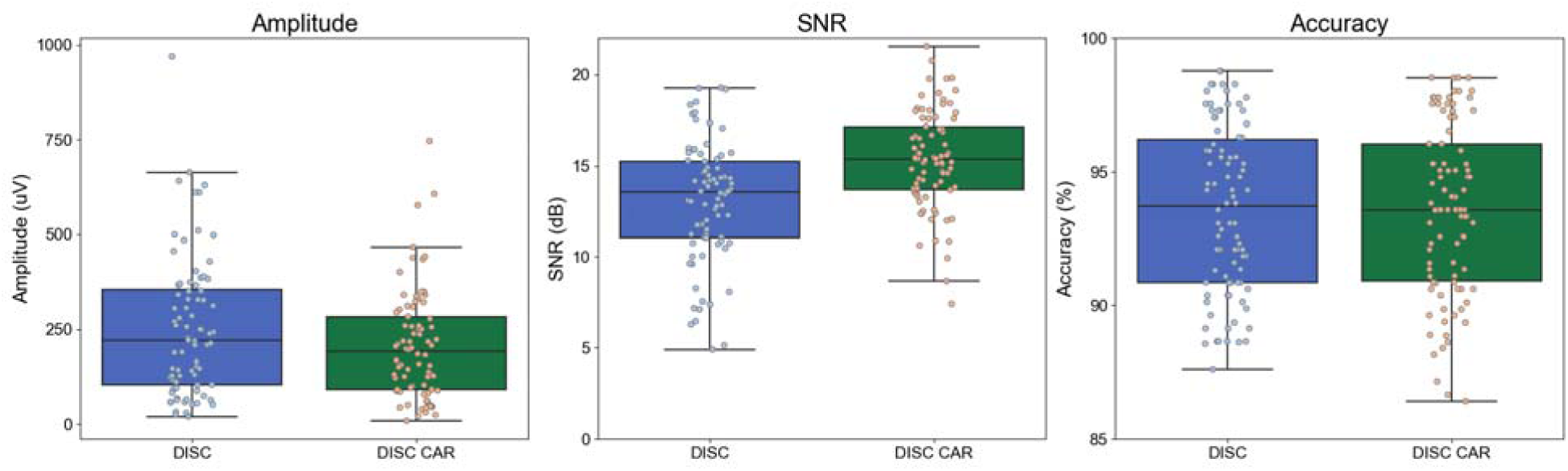
*In vivo* performance of common average referencing (CAR). (N=9 subjects, 9 whiskers each). Amplitude and SNR data points represent the average by whisker and subject. Each accuracy value is the 10-fold cross-validation for the 9-whisker model by subject. Given noise was low due to shielding from a Faraday cage, the advantage of CAR is negligible. These results will vary considerably given environmental factors.

**Suppl. Fig. 6.**
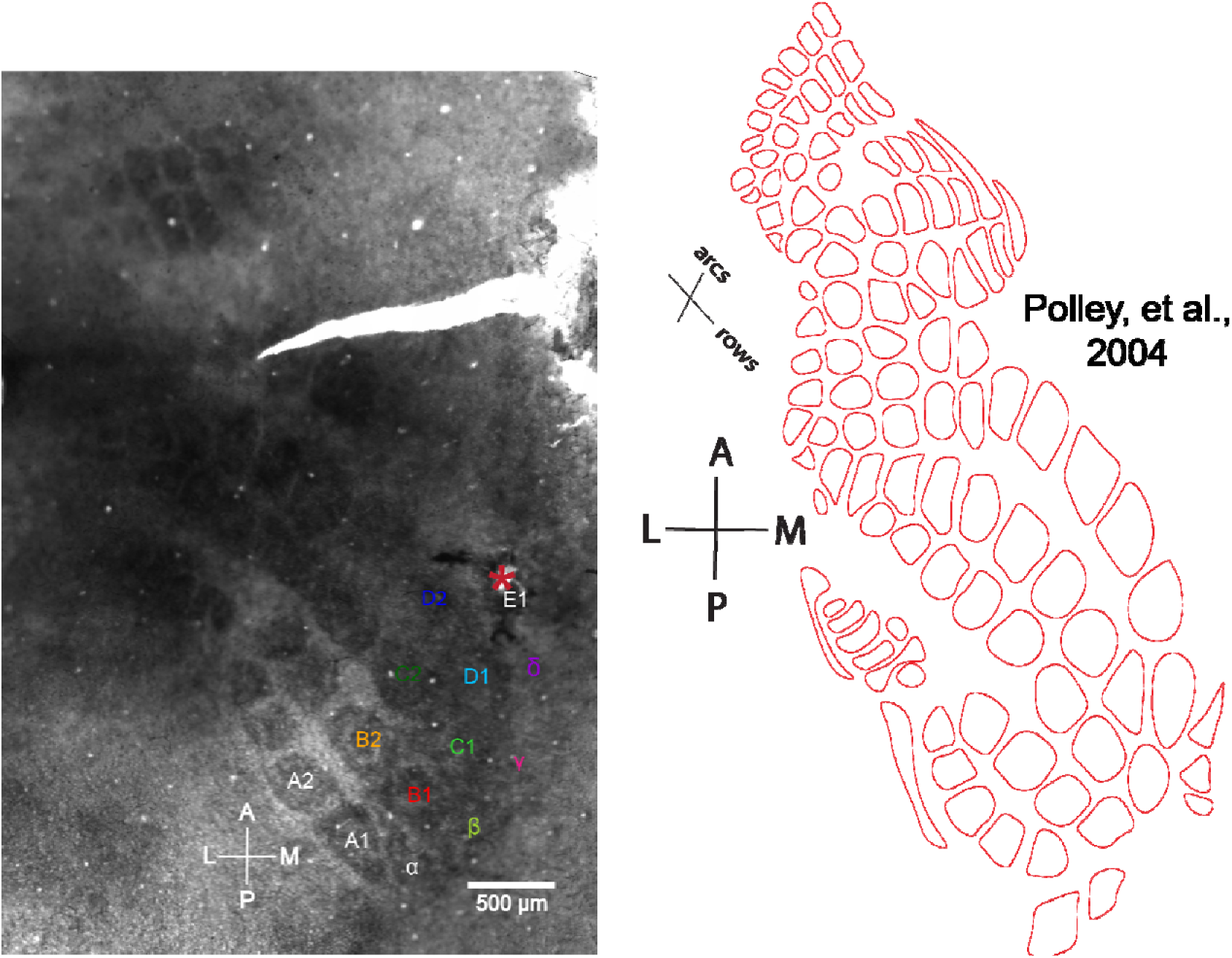
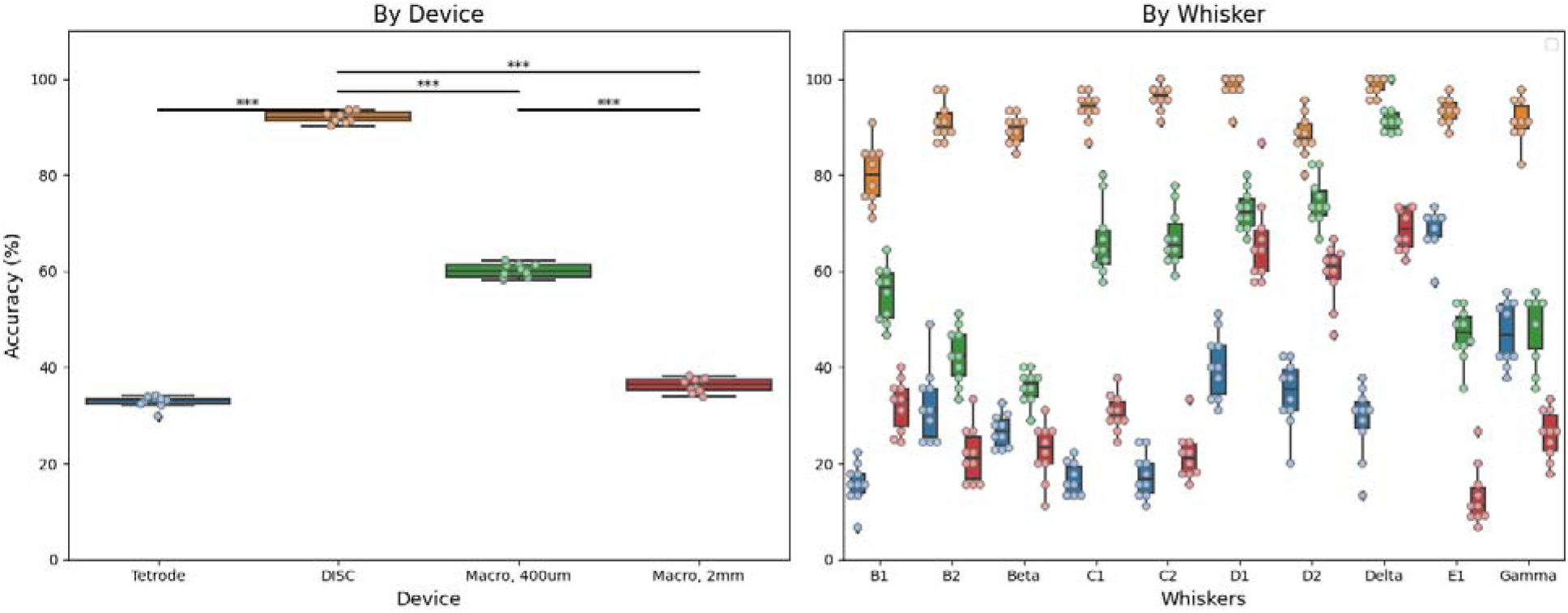
Subject S6 histology and accuracy by whisker. The position of the implant was barrel E1. A useful reference for identifying barrels in rats is Polley et al. ^2^. Further evidence of implanting in E1 was the tetrode amplitude, which was maximum at E1 (data not shown). DISC performed well in classification across the whiskers despite a maximum distance of 1.45 mm. Also promising is the overall performance even though most sources were positioned at roughly the same angle from DISC.

**Suppl. Fig. 7.**
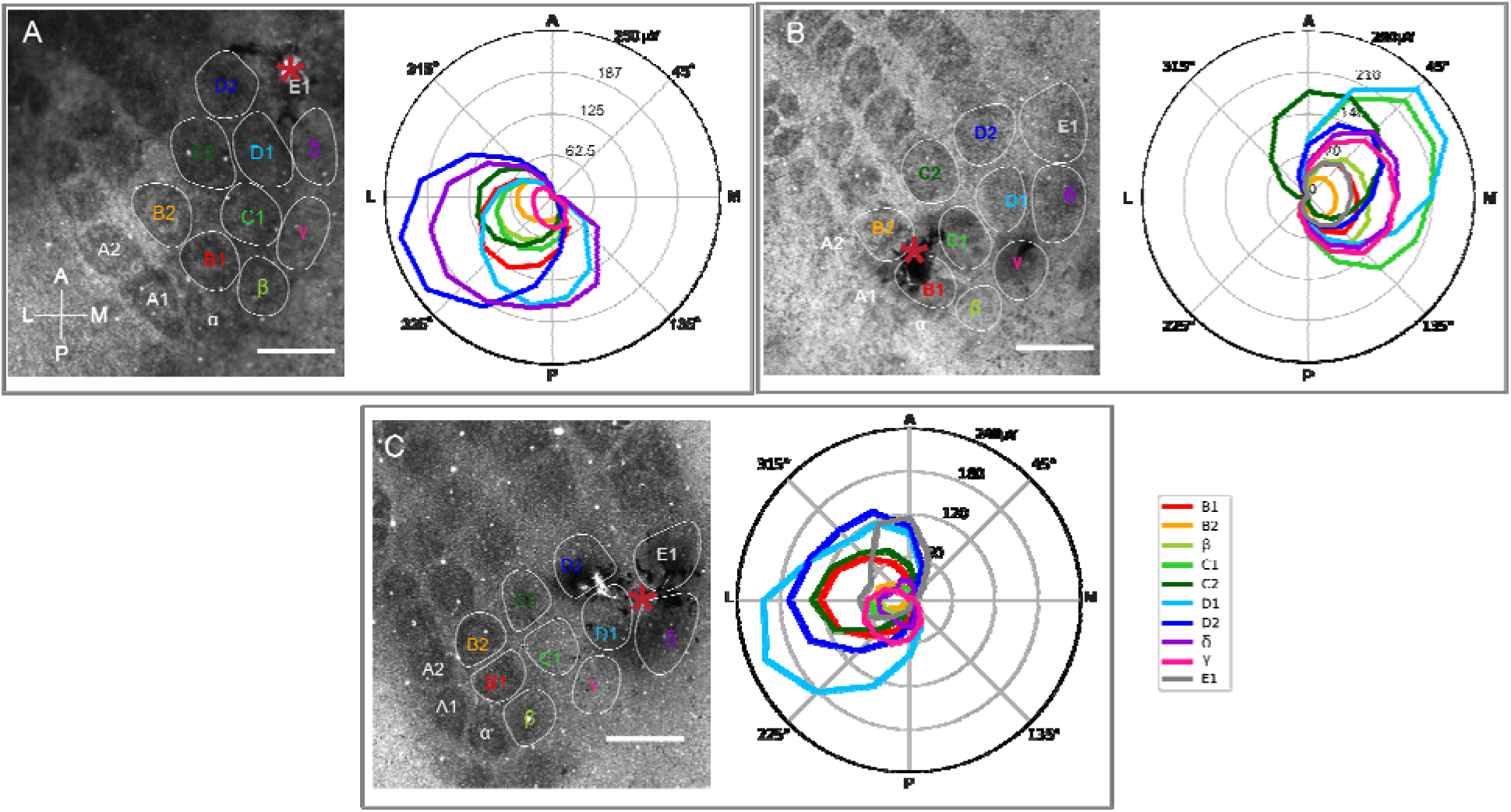
Subject S6 (A), S8 (B), S9 (C) histology and directional curves. The results demonstrate close agreement with histology and the individual whisker response. Directional curve taken from layer V electrode ring.

**Suppl. Fig. 8.**
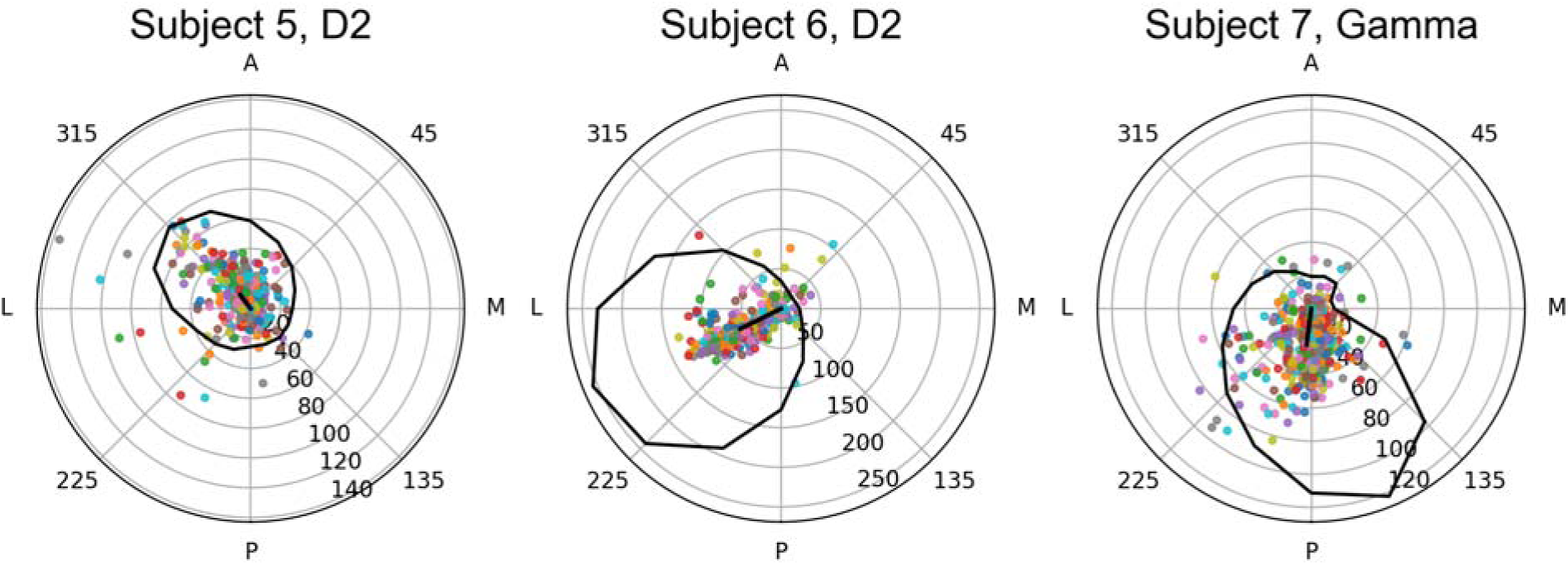
Resultant vector variance. Each colored dot represents the resultant vector for 450 trials for these example whiskers and subjects. The mean resultant vector (black line) and average direction curve illustrate the trend of the data (black closed circle, minimum value subtracted to improve contrast). Related to the results in Suppl. Table 3, the RV is not by itself a highly accurate predictor for whisker identification (i.e., source separation), but it is a contributing factor.

**Suppl. Fig. 9.**
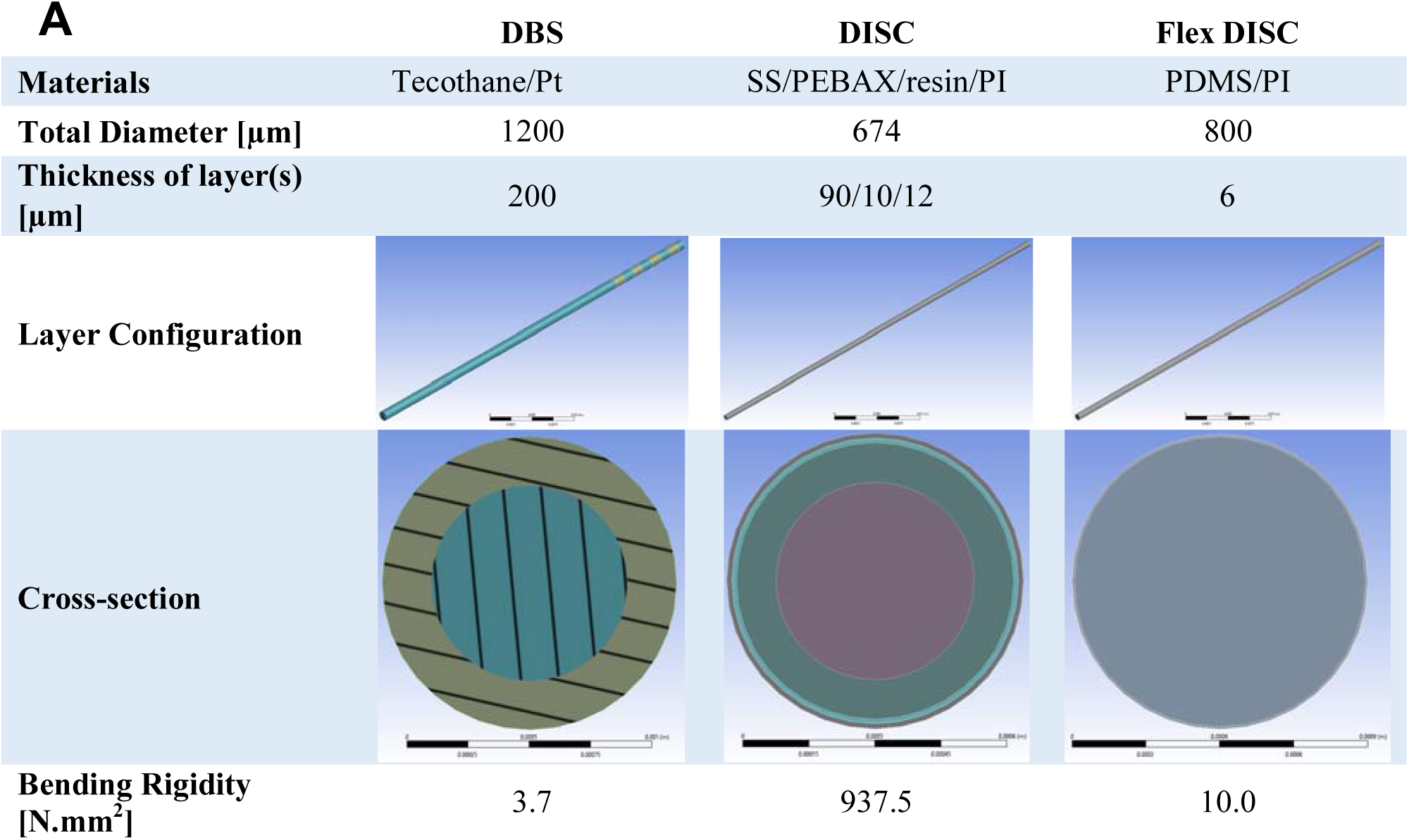
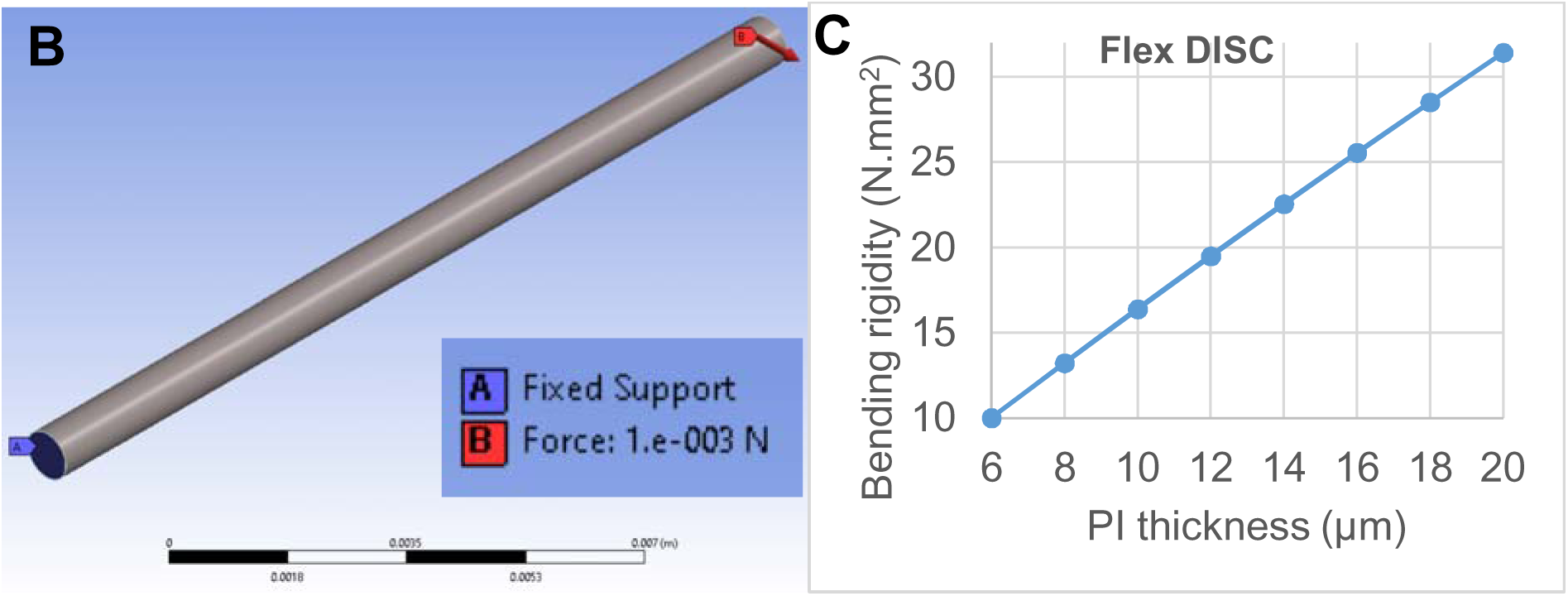
Bending rigidity for DISC and other depth electrode configurations. **A,** Configuration of three devices and their material and geometric properties. **B,** 1 mN force was applied to the tip of 15-mm lead bodies while the other end was fixed using ANSYS mechanical FEM. Bending rigidity (*EI*) is numerically calculated for different electrodes. **C,** The effect of polyimide (PI) thickness on bending rigidity of DISC for an inner cylinder of silicone/PDMS when the total device diameter is held fixed at 800 μm. *Note on Suppl.* Fig. 9: DBS depth arrays are 1.2mm of silicone and have a long history as chronic implants in the human brain, so this was a useful benchmark. The platinum rings (2-mm tall) contributed little to the total rigidity (see Layer configuration). Ideally DISC can be designed to improve the flexibility, but as shown, thin PI film limits our flexibility relative to silicone only. This relationship is almost linear over the chosen range when the total thickness is fixed (C). The predicted thickness of PI required for equivalent rigidity as the modeled DBS device is 1.94 μm, which is feasible but would make assembly and high yield challenging. We also modeled a mesh wrap of PI, but this had a surprisingly small decrease in rigidity (∼20% decrease) and limits the number of channels that can fit on a given layer.

**Suppl. Fig. 10.**
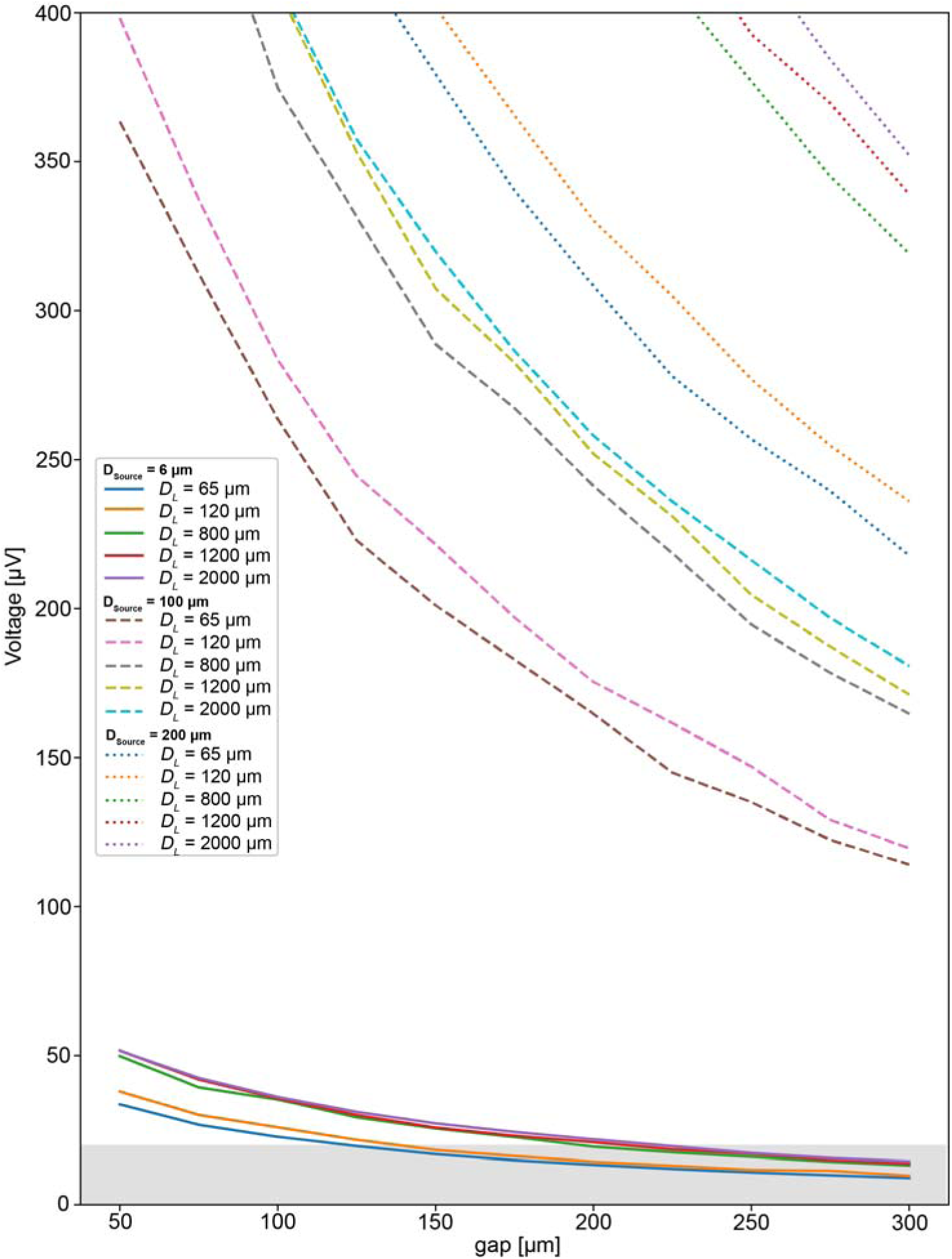
Voltage amplitude predicted for substrate and dipole diameters. The range of a small dipole increases asymptotically with substrate diameter. For example, the 20-μV detection limit of a 6-μm source using a narrow substrate (Ø: 65 μm) is 125 µm. The 800-μm Ø substrate has a predicted range of 195 µm, and 1.2 mm Ø substrate increases to a 205 µm range. *Note on Suppl.* Fig. 10: Two papers from the University of Iowa demonstrate recording in humans with putative single units (N=6 patients)^3^ and multi-unit recordings in humans (N=3 patients)^4^ despite the large diameter of their device (1.2 mm). The quality of their recordings improved on day 5 and afterward. Our modeling of edema vs encapsulation (Suppl. Fig. 10) is consistent with their results. In the latter study, 109 multi-units were sorted using MountainSort across 3 patients, each with 14 high impedance electrodes. This is an average yield of 2.6 distinct multi-units per electrode, which is surprisingly competitive with much smaller intracortical arrays. The literature consistently predicts increased tissue damage with larger diameter devices, but it is worth studying whether the extended range of larger devices will at least mitigate part of the resulting neuronal loss. Biostability, stiffness, and tethering are also expected to be important factors in such a future study.

**Suppl. Fig. 11.**
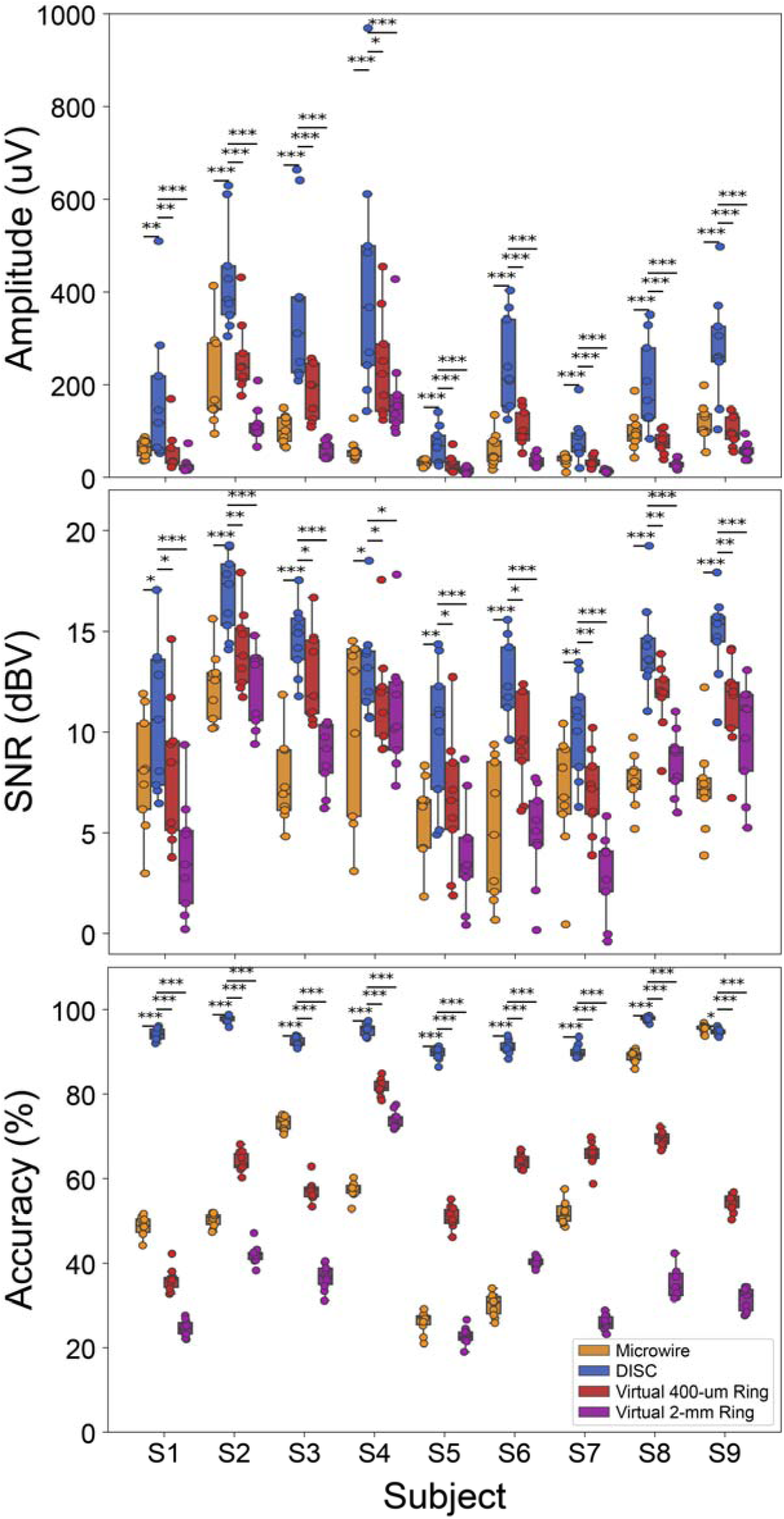
LFP Amplitude, SNR and Accuracy for each device by subject. Improved accuracy of DISC over other device types can be independently tested and shown for each subject. Only S9 failed to show a significant improvement using DISC, where high fidelity oscillations were present post-stimulation on tetrodes and DISC.

Suppl. Movie 1: MP4 of 3D voltage contours for a growing electrode size. *(separate file)*

Suppl. Movie 2: MP4 of 3D voltage contours for various source to device configurations *(separate file)*

## Notes

### Competing Interest Statement

The authors have declared no competing interest.

### Summary of Updates

Version 2 (Oct 2021): Added the final subjects and more supplemental figures. Version 3 (Jan 2022): Added Extended Data figures and several new figures in the supplemental.

